# Structural and functional impact by SARS-CoV-2 Omicron spike mutations

**DOI:** 10.1101/2022.01.11.475922

**Authors:** Jun Zhang, Yongfei Cai, Christy L. Lavine, Hanqin Peng, Haisun Zhu, Krishna Anand, Pei Tong, Avneesh Gautam, Megan L. Mayer, Sophia Rits-Volloch, Shaowei Wang, Piotr Sliz, Duane R. Wesemann, Wei Yang, Michael S. Seaman, Jianming Lu, Tianshu Xiao, Bing Chen

## Abstract

The Omicron variant of severe acute respiratory syndrome coronavirus 2 (SARS-CoV-2), bearing an unusually high number of mutations, has become a dominant strain in many countries within several weeks. We report here structural, functional and antigenic properties of its full-length spike (S) protein with a native sequence in comparison with those of previously prevalent variants. Omicron S requires a substantially higher level of host receptor ACE2 for efficient membrane fusion than other variants, possibly explaining its unexpected cellular tropism. Mutations not only remodel the antigenic structure of the N-terminal domain of the S protein, but also alter the surface of the receptor-binding domain in a way not seen in other variants, consistent with its remarkable resistance to neutralizing antibodies. These results suggest that Omicron S has acquired an extraordinary ability to evade host immunity by excessive mutations, which also compromise its fusogenic capability.

## Introduction

Multiple waves of COVID-19 cases have been driven primarily by emergence of new variants of severe acute respiratory syndrome coronavirus 2 (SARS-CoV-2) since its initial outbreak. The latest surge of new infections is caused by the Omicron variant (*1*), which has acquired an unusually large number of mutations, particularly in its spike (S) protein (~40 residue changes versus 10 on average in all the previous dominant variants; ref (*2*)). This variant, first detected in South Africa and Botswana (also known as lineage B.1.1.529), was designated as a variant of concern (VOC) by the World Health Organization in two days, and has outcompeted the previously most contagious variant - Delta-in many countries within 2-3 weeks. Omicron appears to be much more transmissible than all other variants, as manifested by the sharp increase of new cases in different places (*3, 4*) and by early evidence showing that this variant replicates ~70 times faster than previous variants in the bronchi tissue (but >10 times less efficient in lung tissue; ref(*5*)). The variant appears to cause only attenuated infection and lung disease in rodents (*6*), consistent with the earlier reports that most Omicron cases are generally mild in humans (*7, 8*). Not surprisingly, it has a remarkable ability to evade host immunity generated by either vaccination of the first-generation vaccines or natural infection from previous variants (*9-13*), in agreement with a large volume of breakthrough infection and reinfection cases (*14–17*). It is critical to understand the molecular mechanisms of these extraordinary behaviors of the Omicron variant to tailor effective strategies for controlling its spread.

SARS-CoV-2 is an enveloped positive-stranded RNA virus that can infect a host cell after the virus-encoded trimeric spike (S) protein engages the cellular receptor angiotensin converting enzyme 2 (ACE2) and induces fusion of the viral and target cell membranes. The S protein is first synthesized as a single-chain precursor, and subsequently cleaved by a host furin-like protease into the receptor-binding fragment S1 and the fusion fragment S2 (Fig. S1; ref(*18*)); three copies of each fragment form a noncovalently-associated complex as the mature viral spike. After binding to ACE2 on the host cell surface, the S protein is further cleaved in S2 (S2’ site; Fig. S1) by an another cellular protease, either TMPRSS2 or cathepsins B and L (CatB/L) (*19*), prompting dissociation of S1 and a series of conformational changes in S2. The large structural rearrangements of S2 drive the fusion of viral and cell membranes, thereby delivering the viral genome into the host cell to initiate infection (*20–22*). S1 contains four domains - NTD (N-terminal domain), RBD (receptor-binding domain), and two CTDs (C-terminal domains), and three copies of S1 wrap around the central helical-bundle structure formed by the prefusion S2 trimer on the surface of virion. The RBD fluctuates between a “down” conformation for a receptor-inaccessible state, or an “up” conformation for a receptor-accessible state (*23*), effectively engaging the receptor while protecting the receptor binding site from the host immune system (*23, 24*).

Several structural studies using the stabilized, soluble ectodomain constructs of the Omicron S protein have shown how its mutations modify the antigenic surfaces and how the mutated RBD interacts with ACE2 and monoclonal antibodies (*25–30*). Additional analyses suggest that the function of the Omicron spike is somehow attenuated in cell culture (*31, 32*). In this work, we have characterized the full-length S protein of the Omicron variant and determined its structures with a native sequence by cryogenic electron microscopy (cryo-EM). Comparison of the structure, function and antigenicity of the Omicron S with those of the previously characterized variants (*33*) has given molecular insights into the latest fast-spreading form of SARS-CoV-2.

## Results

### Omicron S requires a higher level of ACE2 than other variants for efficient membrane fusion

To characterize the full-length S protein derived from a natural isolate of Omicron variant (hCoV-19/South Africa/CERI-KRISP-K032233/2021) (Fig. S1), we transfected HEK293 cells with an Omicron S expression construct and compared its membrane fusion activity with that of the full-length S constructs from previous variants, including G614, Alpha, Beta and Delta (*33–35*). All S proteins expressed at comparable levels with the Omicron protein cleaved slightly less between S1 and S2 at 24 hours posttransfection (Fig. S2A), suggesting that the two mutations (N679K and P681H) in Omicron near the furin cleavage site did not enhance the S protein processing. Under our standard assay conditions (*36*), the cells producing these S proteins fused efficiently with ACE2-expressing cells, except that the fusion activity of Omicron S was slightly less than other S proteins (Fig. S2B).

To further test whether Omicron S could induce membrane fusion more efficiently than other variants to account for its rapid spread, we carried out a time-course experiment with our standard cell-cell fusion assay with both S and ACE2 transfected at saturating levels (Fig. 1A; ref (*36*)). As shown previously, there was no significant differences in the fusion activity among all other variants (*33, 35*), but the Omicron S again showed a slightly lower activity throughout the time period tested (Fig. 1A). When using HEK293 cells, which express a minimal level of endogenous ACE2, as the target cells, all other variants (particularly Delta) had significant fusion activities at later time points, while the Omicron S remain inactive (Fig. 1B), suggesting that Omicron S failed to use a low level of ACE to enter host cells. We next tested HEK293 cells transfected with different amounts of ACE2. As shown in Fig 1C, all variants responded to an increased level of ACE2, but the Omicron S lagged behind the other variants and required an almost 10 times higher level of ACE2 to reach a similar fusion activity. The same pattern was also observed when the S-producing cells cotransfected with a furin expression construct and/or the target cells cotransfected with TMPRSS2 (Figs. 1D and S3A, B). We also performed a similar timecourse experiment using murine leukemia virus (MLV)-based pseudoviruses expressing the S constructs with the cytoplasmic tail deleted (*35*). The Delta variant and one of its sublineage variant AY.4 infect the ACE-expressing target cells much more rapidly in the early time period than did any other variant, consistent with our previous results (*35*). The Omicron pseudoviruses, produced either in the absence or in the presence of overexpression of furin, showed faster kinetics of infection than G614, but still substantially slower than the Delta variants in the early time period (Fig. S3C).

**Figure 1.**
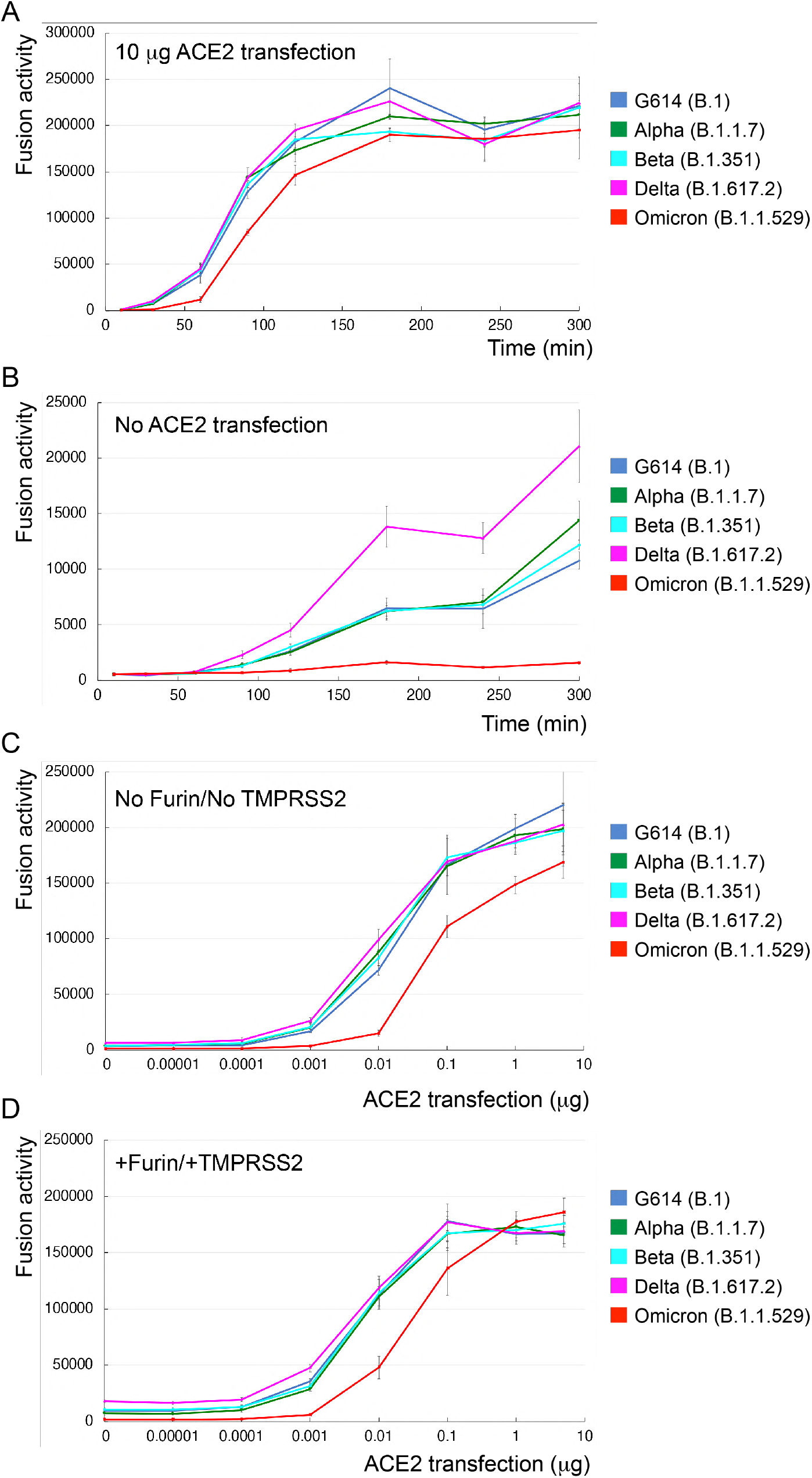
Requirement of higher levels of ACE2 for efficient membrane fusion by the Omicron spike. (**A**) Time-course of cell-cell fusion mediated by various full-length S proteins, as indicated, with the target HEK293 cells transfected with 10 μg ACE2. (**B**) Time-course of cell-cell fusion mediated by various full-length S proteins, as indicated, using HEK293 cells without exogenous ACE2. (**C**) Cell-cell fusion mediated by various full-length S proteins with HEK293 cells transfected with various levels (0-5 μg) of the ACE2 expression construct. (**D**) Cell-cell fusion mediated by various full-length S proteins expressed in HEK293 cells cotransfected with 5 μg furin expression construct and the ACE2-expressing target cells cotransfected with 5 μg TMPRSS2 expression construct. The experiments were repeated at least twice with independent samples giving similar results.

These results together suggest that the Omicron S requires higher levels of the receptor ACE2 for efficient membrane fusion and that it has not gained any obvious advantages in its fusogenecity, as compared to other prevalent variants.

### Biochemical and antigenic properties of the intact Omicron S protein

To produce the full-length Omicron S protein without any stabilizing modifications, we used a C-terminal strep-tagged construct for expression (Fig. S4A), and purified the protein under the identical conditions established for producing other intact S proteins (*33, 34, 36*). By gel-filtration chromatography, the purified Wuhan-Hu-1 S protein can be resolved in three distinct peaks, corresponding to the prefusion S trimer, postfusion S2 trimer and dissociated S1 monomer, respectively, while the G614 S protein elutes as a single peak of the prefusion trimer with no obvious dissociated S1 and S2 (Fig. S4B; ref (*34, 36*)). Although the Omicron protein also eluted in one major peak corresponding to the prefusion trimer, there was a significant amount of aggregate on the leading side and also a shoulder on the trailing side, suggesting that the Omicron protein is less stable than the G614 trimer (Fig. S4B). SDS-PAGE analysis showed that a large fraction of the protein remained uncleaved at the time when the transfected cells were harvested ~84 hours posttransfection (Fig. S4B), indicating that the furin cleavage for the Omicron variant is indeed not very efficient, consistent with the observations reported by others (*31, 32*). These data indicate that the two mutations in Omicron S near the furin site have reduced the proteolytic cleavage, which has been shown to be critical for enhanced viral infectivity and pathogenesis (*37, 38*).

To analyze receptor-binding and antigenic properties of the prefusion Omicron S trimer, we compared its binding to soluble ACE2 proteins and S-directed monoclonal antibodies with that of the G614 trimer by bio-layer interferometry (BLI). The selected antibodies, isolated from COVID-19 convalescent individuals and also used for characterizing previous variants (*33, 35*), target distinct epitopic regions on the S trimer, as defined by competing groups designated RBD-1, RBD-2, RBD-3, NTD-1, NTD-2 and S2 (Fig. S5A; ref(*39*)). Those from the NTD-2 and S2 groups are largely non-neutralizing. The Omicron variant bound substantially more strongly to the receptor than did the G614 trimer, regardless of the ACE2 oligomeric state (Figs. 2 and S5B; Table S1), suggesting that the mutations in the RBD of Omicron S have increased receptor binding affinity and could in principle enhance the Omicron infectivity.

**Figure 2.**
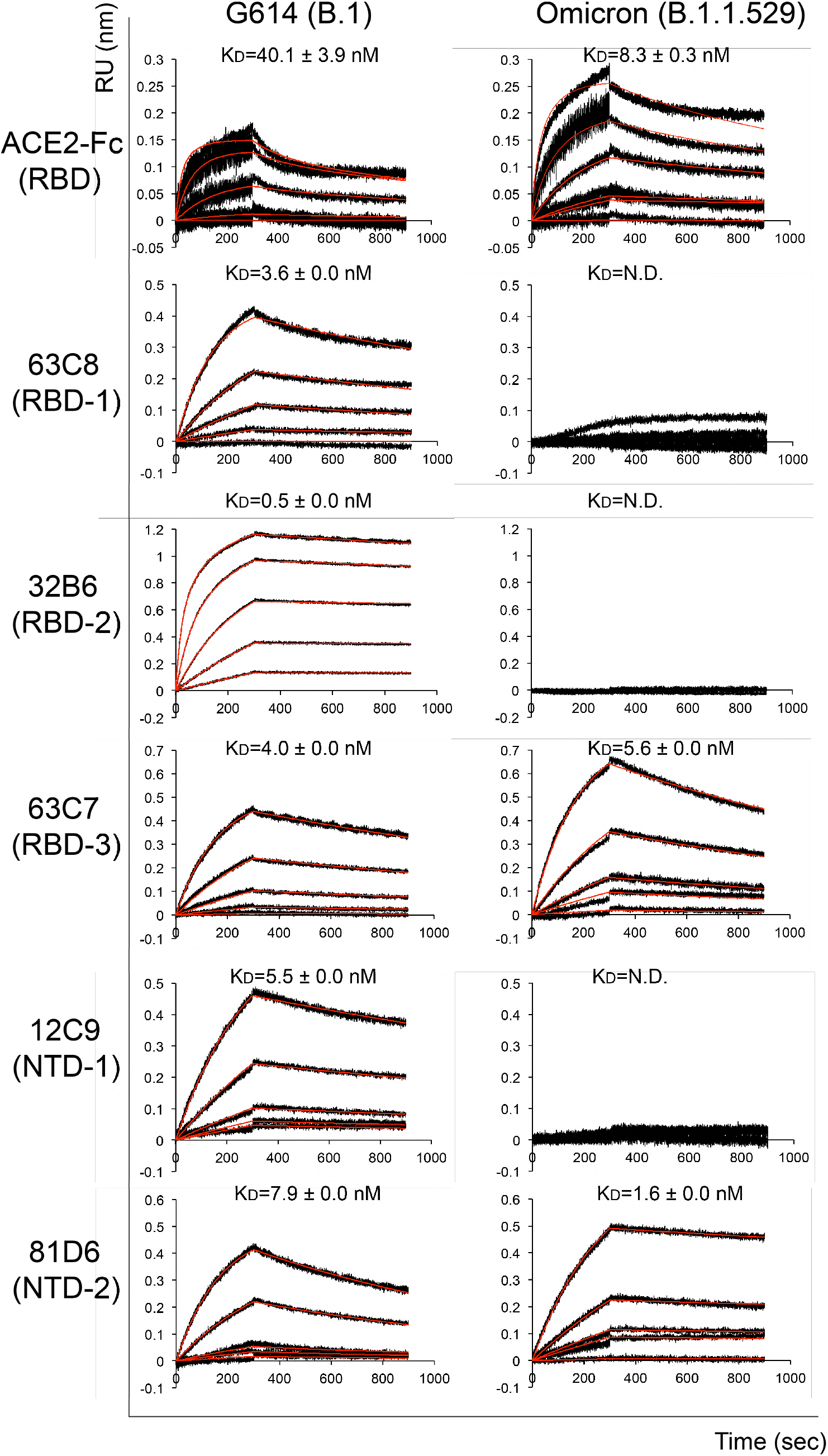
Antigenic properties of the purified full-length Omicron S protein. Bio-layer interferometry (BLI) analysis of the association of prefusion S trimers derived from the G614 “parent” strain (B.1) and Omicron (B.1.1.529) variant with a soluble dimeric ACE2 construct and with a panel of antibodies representing five epitopic regions on the RBD and NTD (see Fig. S5A and ref(*39*)). For ACE2 binding, purified ACE2 protein was immobilized to AR2G biosensors and dipped into the wells containing each purified S proteins at various concentrations. For antibody binding, various antibodies were immobilized to AHC biosensors and dipped into the wells containing each purified S protein at different concentrations. Binding kinetics were evaluated using a 1:1 Langmuir model except for dimeric ACE2 and antibody G32B6 targeting the RBD-2, which were analyzed by a bivalent binding model. The sensorgrams are in black and the fits in red. Binding constants highlighted by underlines were estimated by steady-state analysis as described in the Methods. RU, response unit. Binding constants are also summarized here and in Table S1. N.D., not determined. All experiments were repeated at least twice with essentially identical results.

The selected monoclonal antibodies bind the G614 S trimer with reasonable affinities, as documented previously (*33, 35*). The Omicron variant completely lost binding to both the NTD-1 antibodies, 12C9 and C83B6, both the RBD-2 antibodies, G32B6 and C12A2, as well as the non-neutralizing anti-S2 antibody C163E6, and showed barely detectable binding to the RBD-1 antibody 63C8 (Figs. 2 and S5B; Table S1). Its affinities for the RBD-3 antibody C63C7 and the non-neutralizing NTD-2 antibody C81D6 were the same or even higher than those of the G614 trimer. The BLI data were also largely confirmed by the binding results with the membrane-bound S trimers measured by flow cytometry (Fig. S6).

We also measured the neutralization potency of these antibodies and of a designed trimeric ACE2 (*40*) against infection by the Omicron variant in an HIV-based pseudovirus assay. The Omicron variant was completely resistant to almost all selected antibodies, except for the RBD-3 antibody C63C7 that neutralized very weakly (Table S2), in agreement with their binding affinity for the membrane-bound or purified S proteins, as well as the results reported by others (*9–13*). The two non-neutralizing antibodies, C81D6 and C163E6, did not neutralize the pseudoviruses, as expected. In contrast, the trimeric ACE2 is significantly more potent against the Omicron variant than the G614, consistent with the increased receptor binding by Omicron.

### Overall structure of the full-length S trimer of the Omicron variant

We next determined the cryo-EM structures of the full-length Omicron S trimer with the unmodified sequence. Cryo-EM images were acquired on a Titan Krios electron microscope equipped with a Gatan K3 direct electron detector. We used crYOLO (*41*) for particle picking, and RELION (*42*) for two-dimensional (2D) classification, three dimensional (3D) classification and refinement (Figs. S7 and S8). 3D classification gave three distinct classes for the Omicron S trimer, representing the closed, three-RBD-down prefusion conformation and a one-RBD-up conformation, as well as an RBD-intermediate conformation that was also found for the G614 trimer (*34*). These classes were further refined to 3.1-4.3Å resolution and we modeled the closed and one-RBD-up conformations (Fig. S7-S9; Table S3).

Similar to other variants, there are no major differences in the overall architecture between the full-length Omicron S protein and the G614 S trimer in the corresponding conformation (Fig. 3A and 3B; ref(*34*)). In the closed, three-RBD-down conformation, the NTD, RBD, CTD1 and CTD2 of S1 wrap around the S2 trimer. In the one-RBD-up conformation, the central helical core structure of S2 was preserved despite the flip-up movement of the RBD, which opened up the S1 trimer by shifting two adjacent NTDs away from the three-fold axis of the trimer. Noticeably, additional residues near the furin cleavage site at the S1/S2 boundary (residues 682-685), largely disordered in the previous variants (*33–36*), became structured probably due to the N679K mutation since the sidechain of Lys679 could make some contacts with the nearby CTD-2 (Fig. S10). The reduced flexibility of the cleavage site could slow down its docking into the furin active site, thereby decreasing the cleavage efficiency in the Omicron S. The nearby mutation H655Y, not part of the furin site and also present in the Gamma variant, did not lead to any obvious structural changes (Fig. S10), but changing its interaction with Phe643 in the CTD-2 from cation-π or hydrogen-π to π-π stacking could help destabilize the domain to some extent (*43*).

**Figure 3.**
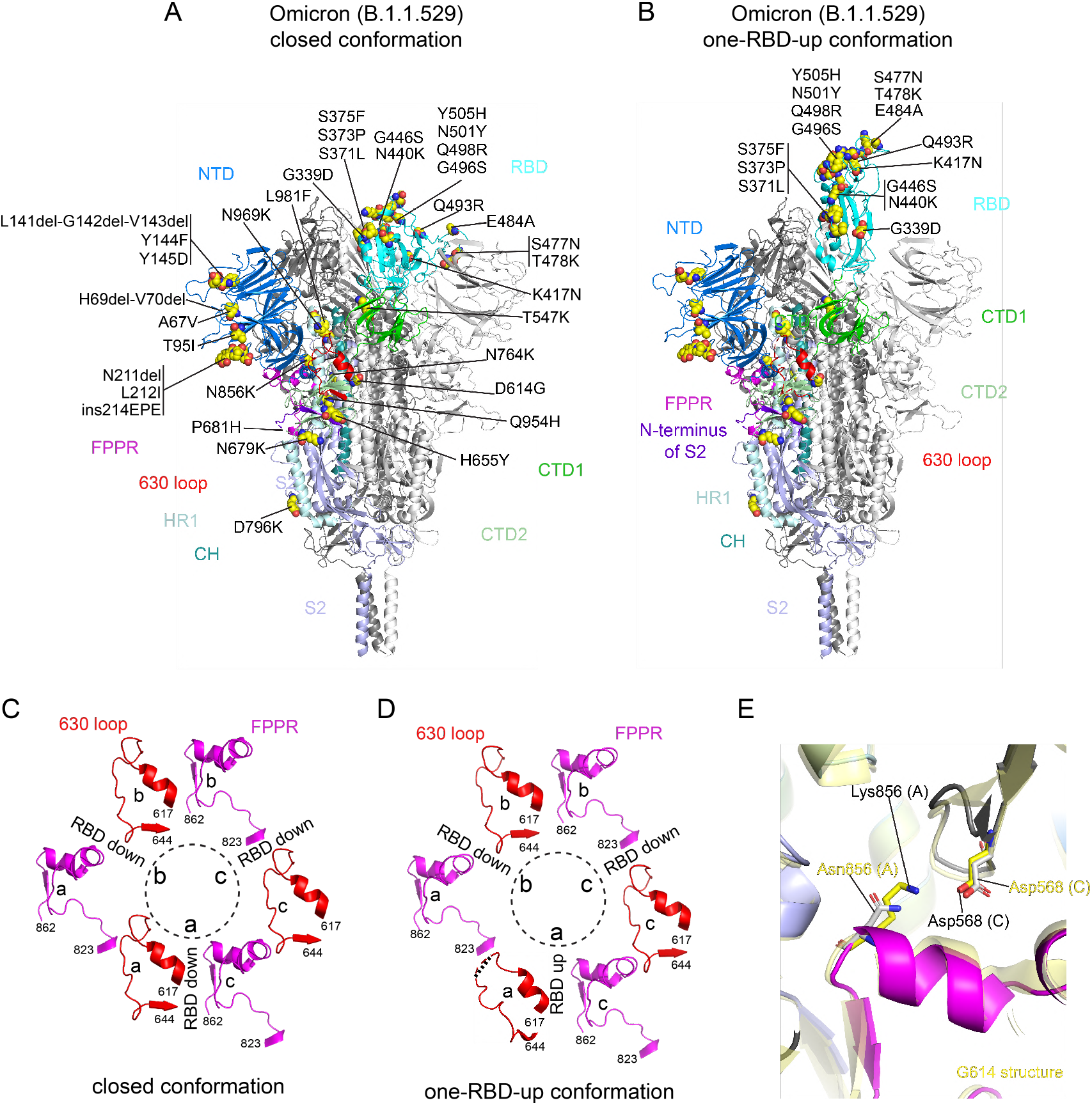
Cryo-EM structures of the full-length Omicron S protein. (**A)** and **(B**) The structures of the closed prefusion conformation and a one-RBD-up conformation of the Omicron S trimer, respectively, are shown in ribbon diagram with one protomer colored as NTD in blue, RBD in cyan, CTD1 in green, CTD2 in light green, S2 in light blue, the 630 loop in red, FPPR in magenta, HR1 in light blue, CH in teal and the N-terminal segment of S2 in purple. All mutations in the Omicron variant, as compared to the original virus (Wuhan-Hu-1), are highlighted in sphere model. (**C**) and (**D**) Structures, in the Omicron closed and one-RBD-up conformations, respectively, of segments (residues 617-644) containing the 630 loop (red) and segments (residues 823-862) containing the FPPR (magenta) from each of the three protomers (a, b and c). The position of each RBD is indicated. Dashed lines indicate gaps in the chain trace (disordered loops). (**E**) Superposition of the structure of the Omicron S trimer in various colors with that of the G614 trimer in yellow aligned by S2, showing the region near the mutation N856K.

Our previous studies indicate that the FPPR (fusion peptide proximal region; residues 828 to 853) and 630 loop (residues 620 to 640) are control elements and their positions may modulate the RBD movement and thereby the kinetics of the S structural rearrangements (*34, 36, 44*). The RBD up movement apparently pushes the FPPR and 630 loop out of their original positions in the closed conformation, making them invisible in cryo-EM maps. In the three-RBD-down conformation of the Omicron trimer (Fig. 3C), the configurations of the FPPR and 630 loop are identical to the distribution observed in the G614 trimer and they are all structured. In the one-RBD-up conformation of the G614 trimer, only one the FPPR and 630-loop pair is ordered (*34*). Unexpectedly, all three FPPRs and two 630 loops in the one-RBD-up Omicron trimer remain structured and largely maintain their positions found in the RBD-down conformation; only the 630 loop immediately next to the RBD in the up conformation became partially disordered (Fig. 3D). The mutation N856K has apparently created a new salt bridge between the FPPR (Lys856) and the CTD-1 (Asp568) of the neighboring protomer (Fig. 3D), stabilizing the FPPR even when the RBD above it adopted an up-conformation. As a result, the flipped RBD together with the CTD-1 from the same protomer had to move up by 3-4Å in order to avoid clashes with the FPPR underneath, probably also leaving more space for the 630 loop (Fig. S11).

Density for the N-linked glycan at residue Asn343 in the RBD has rotated up almost by 90° in the Omicron map, as compared to that of the G614 and Delta trimers (Fig. S12A). The distal end of the glycan was found to make contacts with the neighboring RBD, helping clamp down the three RBDs in the closed conformation of the Delta trimer. Shifting away the N343-glycan would weaken the packing between the neighboring RBDs, at least partially accounting for the slight outward movement of all three RBDs in the Omicron trimer (Fig. S12B).

### Structural consequences of mutations in the Omicron variant

The overall structure of the RBD has changed little between the Omicron and G614 trimers (Fig. 4A), except for a small shift of a short helix (residues 365-371; Fig. 4B). The surface along the RBM (receptor binding motif) exposed in the closed conformation, which include the ACE2 binding interface and many antibody epitopes from the RBD-1 and RBD-2 groups (Fig. S4A), has been modified by the large number of mutations, however. N501Y is present in the previous variants and has been shown to enhance ACE2 affinity by making additional contacts (*45*). New mutations Q493R and Q498R can create new salt bridges with residues from ACE2, as confirmed by recent studies (*25, 27–29*). K417N and E484A (or some forms) are also found in other variants and they may reduce the ACE2 affinity because of loss of ionic interactions with the receptor (*33, 46*). In combination, these mutations may be responsible for the observed increase in the receptor binding reported here and by others (*25, 27, 29*). Three mutations, S477N, T478K and E484A, in the tip of the RBM, possibly together with K417N, can probably account for loss of binding and neutralization of the Omicron variant by antibodies that target the RBD-2 epitopes. Other mutations, such as, G339, N440K, G446S, Q493R, G486S, Q498R, Y505H, all aligned near the RBD-1 epitopes, are probably responsible for resistance to the antibodies of this group. There are few mutations near the RBD-3 epitopes (the “so-called cryptic site”), explaining why C63C7 retained its binding to the Omicron S.

**Figure 4.**
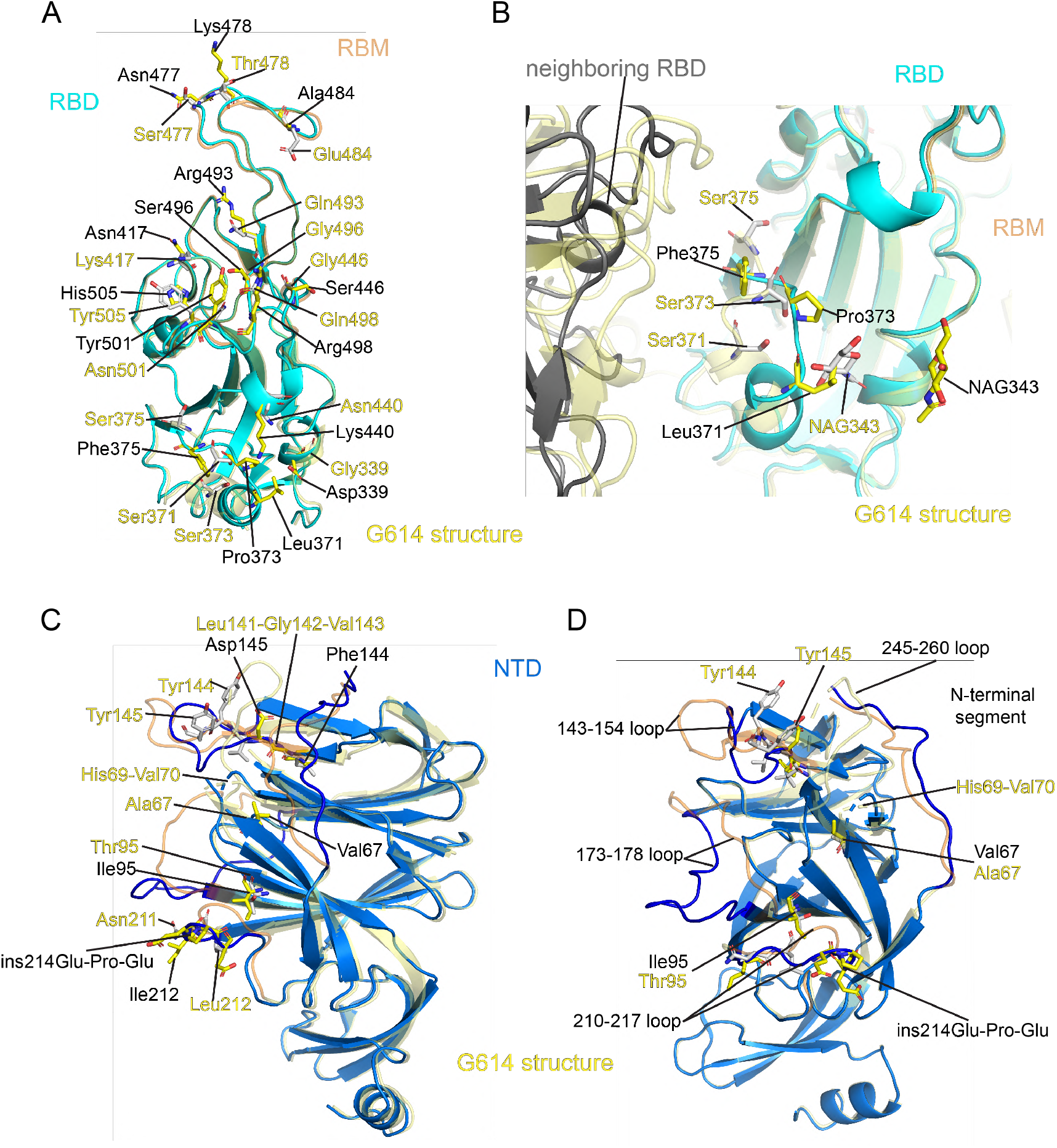
Structural impact of the mutations in the Omicron S. (**A**) Superposition of the RBD structure of the Omicron S trimer in cyan with the RBD of the G614 S trimer in yellow. Locations of all 15 mutations in the RBD are indicated and these residues are shown in stick model. The RBM (receptor binding motif) is colored in orange in the G614 structure. (**B**) A close-up view of the RBD superposition in (A) to show the region near the mutations S371L, S373P and S375F, including part of the neighboring RBD from another protomer. The mutated residues and the N-linked glycans at Asn343 are in stick model. NAG, N-acetylglucosamine. (**C**) and (**D**) Two different views of superposition of the NTD structure of the Omicron S trimer in blue with the NTD of the G614 S trimer in yellow. Locations of mutations A67V, T95I, Y144F, Y145D, L212I, deletions H69del-V70del, L141del-G142del-V143del, N211del and an insertion ins214EPE are indicated and these residues are shown in stick model. The N-terminal segment, 143-154, 173-187, 210-217 and 245-260 loops are rearranged between the two structures and highlighted in darker colors.

The mutation S373P may have rigidified the local polypeptide chain, causing an inward shift of the helix formed by residues 365-371, which in turn led to a large rotation of the N-linked glycan at Asn343 (Figs. 4B and S12A). Both the shift of the helix and the glycan would lose contacts with the RBD from the neighboring protomer and relax the inter-RBD packing, although these interactions may not be functionally critical because the density of the distal end of N343-glycan in the G614 map is weaker than that in the Delta trimer (Fig. S12A)

When the structures of the Omicron and G614 S trimers in the closed conformation were superposed by the S2 region, the most prominent differences were in the NTD (Fig. S12B), which contains five point mutations (A67V, T95I, Y144F, Y145D and L212I), three deletions (H69del-V70del, L141del-G142del-V143del and N211del) and one three-residue insertion (ins214EPE). When the two NTDs were aligned (Fig. 4C and 4D), we could see that these mutations had reconfigured the N-terminal segment and almost all the surface-exposed loops, including the 143-154, 173-187, 210-217 and 245-260 loops. Many of the loops form important parts of the neutralizing epitopes in the NTD (*47*). Thus, these structural changes, substantially different from those found in the previous variants, have once again drastically altered the antigenic surface of the NTD in a unique way, accounting for the loss of binding and neutralization by NTD-1 antibodies (Figs. 2 and S5; Table S2). It is unclear why the affinity by the NTD-2 antibody C81D6 has increased as its epitope has not been defined structurally.

Finally, the rest of the mutations, including N764K, D796Y, Q954H, N969K and L891F, did not lead to any obvious structural changes (Fig. S13A-D), at least in these prefusion conformations, even though some of them are non-conservative changes.

## Discussion

New variants, such as Omicron, may continue to emerge from the worldwide spread of SARS-CoV-2, because the virus has to evolve in order to survive under the selective pressure exerted by increasingly prevalent host immunity from either natural infection or vaccination at the population level. How the Omicron has acquired a much larger number of mutations than all other previous variants is unknown, although origins from immune-compromised individuals or even animals have been suggested (*48, 49*). From our study and those from others (*31, 32*), the Omicron S protein fits the classical description in which an excessively mutated virus has to compromise its fitness in exchange for the ability to evade host immunity, as its fusogenic capability appears to have been attenuated. We cannot rule out the possibility that the Omicron S under certain optimal conditions (e.g., combination of perfect levels of ACE2, furin and TMPRSS2 or other co-receptors present in specific tissues), which have not been tested here, is more fusogenic than all other variants, or that the viral replication is drastically increased by mutations outside of the spike gene. We have indeed observed increased binding of the Omicron S to the ACE2 receptor, but it is puzzling why it consistently shows weaker membrane fusion activity and infectivity in the form of either authentic virus or pseudovirus (*31, 32, 50*). A key finding from our study is that the Omicron S is unique in requiring a much higher level of ACE2 for efficient membrane fusion than all other variants tested. This observation can account for the reduced fusion activity when compared with other variants under the same conditions. It can also explain why Omicron replicates better in the upper respiratory tract than lung as the former has much higher levels of ACE2 (*51, 52*). The transition from the closed to the RBD-up conformation in an S trimer typically requires an order-disorder transition in the FPPR and 630 loop. The structured FPPRs and 630 loops found in the Omicron trimer, in particular, with the one-RBD-up conformation, would substantially slow down the transition to two-RBD-up or three-RBD-up conformations, which may be the key trigger for S1 dissociation and S2 refolding required for driving the membrane fusion process. The added kinetic barriers by the structured FPPR and 630 loop would need a higher concentration of ACE2 to overcome.

If the Omicron S is indeed compromised in its fusogenicity, how can we explain the unprecedented spreading of this variant? First, the extraordinary ability of Omicron to evade host immunity would drastically expand the susceptible population and it would be difficult to break the transmission chain until the immunity against Omicron can be rebuilt again by vaccination or infection. A recent study demonstrated that the household secondary attack rate by Omicron is similar to that by Delta among unvaccinated individuals, but substantially higher in those fully vaccinated and booster-vaccinated (*53*), suggesting that the rapid spread by Omicron may primarily be due to immune evasion rather than heightened transmissibility. Moreover, the vial loads from Omicron infection do not seem to be significantly higher than other variants (*54*). Second, because the Omicron variant causes mainly mild diseases, there might be a greater number of asymptomatic cases than the surges by the previous variants, which could facilitate the spread by the apparently “healthy” individuals. Future studies will be undoubtedly needed to address this important question.

We suggested previously that the RBD is a better target for intervention because the NTD can be remodeled differently by different variants while the overall structure of the RBD has been strictly preserved (*35*). The Omicron structures further support this notion as this variant has also invented yet another new way, including point mutations, deletions and even an insertion, to modify the NTD, while the overall RBD structure has only a small change, even though ~8% of all RBD residues are mutated, but without any deletions or insertions. It appears that the virus must keep the RBD structure intact in order to maintain its ability to engage the receptor and its fitness. Indeed, neutralizing potency of certain anti-RBD antibodies (e.g., Sotrovimab (*55*)) remain unchanged against the Omicron virus, raising the hope that broadly neutralizing therapeutic antibodies and next-generation vaccines can be developed against the RBD of SARS-CoV-2.

## Materials and Methods

### Expression constructs

The gene of a full-length spike (S) protein from Omicron variant (hCoV-19/South Africa/CERI-KRISP-K032233/2021; GISAID accession ID: EPI_ISL_6699757) was assembled from DNA fragments synthesized by Integrated DNA Technologies, Inc. (Coralville, Iowa). The S gene was fused with a C-terminal twin Strep tag (SGGGSAWSHPQFEKGGGSGGGSGGSSAWSHPQFEK) and cloned into a mammalian cell expression vector pCMV-IRES-puro (Codex BioSolutions, Inc, Gaithersburg, MD). The furin and TMPRSS2 expression constructs were purchased from Origene (Rockville, MD, Cat# SC118550 and CAT# SC323858).

### Expression and purification of recombinant proteins

Expression and purification of the full-length S proteins were carried out as previously described (*36*). Briefly, expi293F cells (ThermoFisher Scientific, Waltham, MA) were transiently transfected with the S protein expression constructs. To purify the S protein, the transfected cells were lysed in a solution containing Buffer A (100 mM Tris-HCl, pH 8.0, 150 mM NaCl, 1 mM EDTA) and 1% (w/v) n-dodecyl-β-D-maltopyranoside (DDM) (Anatrace, Inc. Maumee, OH), EDTA-free complete protease inhibitor cocktail (Roche, Basel, Switzerland), and incubated at 4°C for one hour. After a clarifying spin, the supernatant was loaded on a strep-tactin column equilibrated with the lysis buffer. The column was then washed with 50 column volumes of Buffer A and 0.3% DDM, followed by additional washes with 50 column volumes of Buffer A and 0.1% DDM, and with 50 column volumes of Buffer A and 0.02% DDM. The S protein was eluted by Buffer A containing 0.02% DDM and 5 mM d-Desthiobiotin. The protein was further purified by gel filtration chromatography on a Superose 6 10/300 column (GE Healthcare, Chicago, IL) in a buffer containing 25 mM Tris-HCl, pH 7.5, 150 mM NaCl, 0.02% DDM. All RBD proteins were purchased from Sino Biological US Inc (Wayne, PA).

The monomeric ACE2 or dimeric ACE2 proteins were produced as described (*40*). Briefly, Expi293F cells transfected with monomeric ACE2 or dimeric ACE2 expression construct and the supernatant of the cell culture was collected. The monomeric ACE2 protein was purified by affinity chromatography using Ni Sepharose excel (Cytiva Life Sciences, Marlborough, MA), followed by gel filtration chromatography. The dimeric ACE2 protein was purified by GammaBind Plus Sepharose beads (GE Healthcare), followed gel filtration chromatography on a Superdex 200 Increase 10/300 GL column. All the monoclonal antibodies were produced as described (*39*).

### Western blot

Western blot was performed using an anti-SARS-COV-2 S antibody following a protocol described previously (*56*). Briefly, full-length S protein samples were prepared from cell pellets and resolved in 4-15% Mini-Protean TGX gel (Bio-Rad, Hercules, CA) and transferred onto PVDF membranes. Membranes were blocked with 5% skimmed milk in PBS for 1 hour and incubated a SARS-CoV-2 (2019-nCoV) Spike RBD Antibody (Sino Biological Inc., Beijing, China, Cat: 40592-T62) for another hour at room temperature. Alkaline phosphatase conjugated anti-Rabbit IgG (1:5000) (Sigma-Aldrich, St. Louis, MO) was used as a secondary antibody. Proteins were visualized using one-step NBT/BCIP substrates (Promega, Madison, WI).

### Cell-cell fusion assay

The cell-cell fusion assay, based on the α-complementation of E. coli β-galactosidase, was used to measure fusion activity of SARS-CoV2 S proteins, as described (*36*). Briefly, to produce S-expressing cells, HEK293T cells were transfected by polyethylenimine (PEI; 80 μg) with either 5 or 10 μg of the full-length SARS-CoV2 (G614, Alpha, Beta, Delta or Omicron) S construct, as indicated in each specific experiment, and the α fragment of E. coli β-galactosidase construct (10 μg), with/without 5 μg of the fruin expression construct, as well as the empty vector to make up the total DNA amount to 20 μg. To create target cells, the full-length ACE2 construct at various amount (10 pg-10 μg), as indicated in each specific experiment, the ω fragment of E. coli β-galactosidase construct (10 μg), with/without 5 μg of the TMPRSS-2 construct as well as the empty vector were used to transfect HEK293T cells. After incubation at 37°C for 24 hrs, the cells were detached using PBS buffer and resuspended in complete DMEM medium. 50 μl S-expressing cells (1.0×10^6^ cells/ml) were mixed with 50 μl ACE2-expressing target cells (1.0×10^6^ cells/ml) to allow cell-cell fusion to proceed at 37 °C for 2 hrs for our standard assays or from 10 min to 5 hours for the time-course experiments. Cell-cell fusion activity was quantified using a chemiluminescent assay system, Gal-Screen (Applied Biosystems, Foster City, CA), following the standard protocol recommended by the manufacturer. The substrate was added to the cell mixture and allowed to react for 90 min in dark at room temperature. The luminescence signal was recorded with a Synergy Neo plate reader (Biotek, Winooski, VT).

### Binding assay by bio-layer interferometry (BLI)

Binding of monomeric or dimeric ACE2 to the full-length Spike protein of each variant was measured using an Octet RED384 system (ForteBio, Fremont, CA). Briefly, monomeric ACE2 or dimeric ACE2 was immobilized to Amine Reactive 2nd Generation (AR2G) biosensors (ForteBio, Fremont, CA) and dipped in the wells containing the S protein at various concentrations (G614 or Omicron, 0.617-150 nM) for association for 5 minutes, followed by a 10 min dissociation phase in a running buffer (PBS, 0.02% Tween 20, 0.02% DDM, 2 mg/ml BSA). To measure binding of a full-length S protein to monoclonal antibodies, the antibody was immobilized to anti-human IgG Fc Capture (AHC) biosensor (ForteBio, Fremont, CA) following a protocol recommended by the manufacturer. The full-length S protein was diluted using a running buffer (PBS, 0.02% Tween 20, 0.02% DDM, 2 mg/ml BSA) to various concentrations (0.617-50 nM) and transferred to a 96-well plate. The sensors were dipped in the wells containing the S protein solutions for 5 min, followed with a 10 min dissociation phase in the running buffer. Control sensors with no ACE2 or antibody were also dipped in the S protein solutions and the running buffer as references. Recorded sensorgrams with background subtracted from the references were analyzed using the software Octet Data Analysis HT Version 12.0 (ForteBio). Binding kinetics was evaluated using a 1:1 Langmuir model except for dimeric ACE2 and antibodies G32B6 and C12A2, which were analyzed by a bivalent binding model. All K_D_ values for multivalent interactions with antibody IgG or dimeric ACE2 and trimeric S protein are the apparent affinities with avidity effects.

### Flow cytometry

Expi293F cells (ThermoFisher Scientific) were grown in Expi293 expression medium (ThermoFisher Scientific). Cell surface display DNA constructs for the SARS-CoV-2 spike variants together with a plasmid expressing blue fluorescent protein (BFP) were transiently transfected into Expi293F cells using ExpiFectamine 293 reagent (ThermoFisher Scientific) per manufacturer’s instruction. Two days after transfection, the cells were stained with primary antibodies at 10 μg/ml concentration. For antibody staining, an Alexa Fluor 647 conjugated donkey anti-human IgG Fc F(ab’)2 fragment (Jackson ImmunoResearch, West Grove, PA) was used as secondary antibody at 5 μg/ml concentration. Cells were run through an Intellicyt iQue Screener Plus flow cytometer. Cells gated for positive BFP expression were analyzed for antibody and ACE2_615_-foldon T27W binding. The flow cytometry assays were repeated three times with essentially identical results.

### MLV-based pseudovirus assay

Murine Leukemia Virus (MLV) particles (plasmids of the MLV components kindly provided by Dr. Gary Whittaker at Cornell University and Drs. Catherine Chen and Wei Zheng at National Center for Advancing Translational Sciences, National Institutes of Health), pseudotyped with various SARS-CoV-2 S protein constructs, were generated in HEK 293T cells, following a protocol described previously for SARS-CoV (*57, 58*). To enhance incorporation of S protein into the particles, the C-terminal 19 residues in the cytoplasmic tail of each S protein were deleted. To increase the cleavage between S1 and S2, 1.5 μg of the furin expression construct was added into the DNA mixture (20 μg) for MLV particle production. To prepare for infection, 7.5×10^3^ of HEK 293 cells, stably transfected with a full-length human ACE2 expression construct, in 15 μl culture medium were plated into a 384-well white-clear plate coated with poly-D-Lysine to enhance the cell attachment. On day 2, 15 μl of MLV pseudoviruses for each variant were added into each well pre-seeded with HEK293-ACE2 cells. The plate was centrifuged at 114 xg for 5 min at 12°C. After incubation of the pseudoviruses with the cells for a time period (10 min-8 hr), as indicated in the figures, the medium was removed and the cells were washed once with 1xDPBS. 30 μl of fresh medium was added back into each well. The cells were then incubated at 37°C for additional 40 hr. Luciferase activities were measured with Firefly Luciferase Assay Kit (CB-80552-010, Codex BioSolutions Inc).

### HIV-based pseudovirus assay

Neutralizing activity against SARS-CoV-2 pseudovirus was measured using a single-round infection assay in 293T/ACE2 target cells. Pseudotyped virus particles were produced in 293T/17 cells (ATCC) by co-transfection of plasmids encoding codon-optimized SARS-CoV-2 full-length S constructs, packaging plasmid pCMV DR8.2, and luciferase reporter plasmid pHR’ CMV-Luc. G614 S, Omicron S, packaging and luciferase plasmids were kindly provided by Drs. Barney Graham and Tongqing Zhou (Vaccine Research Center, NIH). The 293T cell line stably overexpressing the human ACE2 cell surface receptor protein was kindly provided by Drs. Michael Farzan and Huihui Ma (The Scripps Research Institute). For neutralization assays, serial dilutions of monoclonal antibodies (mAbs) were performed in duplicate followed by addition of pseudovirus. Pooled serum samples from convalescent COVID-19 patients or pre-pandemic normal healthy serum (NHS) were used as positive and negative controls, respectively. Plates were incubated for 1 hour at 37°C followed by addition of 293/ACE2 target cells (1×10^4^/well). Wells containing cells + pseudovirus (without sample) or cells alone acted as positive and negative infection controls, respectively. Assays were harvested on day 3 using Promega BrightGlo luciferase reagent and luminescence detected with a Promega GloMax luminometer. Titers are reported as the concentration of mAb that inhibited 50% or 80% virus infection (IC_50_ and IC_80_ titers, respectively). All neutralization experiments were repeated twice with similar results.

### Cryo-EM sample preparation and data collection

To prepare cryo EM grids, 4.0 μl of the freshly purified full-length Omicron S trimer from the peak fraction in DDM, concentrated to ~3.5 mg/ml was applied to a 1.2/1.3 Quantifoil gold grid (Quantifoil Micro Tools GmbH), which were glow discharged with a PELCO easiGlow™ Glow Discharge Cleaning system (Ted Pella, Inc.) for 60 s at 15 mA in advance. Grids were immediately plunge-frozen in liquid ethane using a Vitrobot Mark IV (ThermoFisher Scientific), and excess protein was blotted away by using grade 595 filter paper (Ted Pella, Inc.) with a blotting time of 4 s, a blotting force of −12 at 4°C with 100% humidity. The grids were first screened for ice thickness and particle distribution. Selected grids were used to acquire images by a Titan Krios transmission electron microscope (ThermoFisher Scientific) operated at 300 keV and equipped with a BioQuantum GIF/K3 direct electron detector. Automated data collection was carried out using SerialEM version 3.8.6 (*59*) at a nominal magnification of 105,000× and the K3 detector in counting mode (calibrated pixel size, 0.83 Å) at an exposure rate of 13.362 electrons per pixel per second. Each movie add a total accumulated electron exposure of ~53.592 e-/Å^2^, fractionated in 50 frames. Data sets were acquired using a defocus range of 0.5-2.2 μm.

### Image processing and 3D reconstructions

Drift correction for cryo-EM images was performed using MotionCor2 (*60*), and contrast transfer function (CTF) was estimated by Gctf (*61*) using motion-corrected sums without dose-weighting. Motion corrected sums with dose-weighting were used for all other image processing. crYOLO (*41*) was used for particle picking and RELION3.0.8 (*42*) for 2D classification, 3D classification and refinement procedure. 3,873,988 particles were extracted from 34,031 images using crYOLO with a trained model, and then subjected to 2D classification, giving 2,091,339 good particles. A low-resolution negative-stain reconstruction of the Wuhan-Hu-1 (D614) sample was low-pass filtered to 40Å resolution and used as an initial model for 3D classification with C1 symmetry. After the first round of 3D classification, a major class with clear structural features was re-extracted with a smaller boxsize and subjected to a second round of 3D classification in C1 symmetry, giving one major class apparently in the closed conformation, and five smaller classes with poor map quality. The major class was re-extracted/unbinned and subjected to one round of 3D auto-refinement, yielding a map at 3.7Å resolution from 210,415 particles. An additional round of signal-subtraction and 3D classification using a mask for the apex region of the S trimer were performed, leading to three distinct groups, one with one class in the closed, RBD-down conformation, the second with two classes in the RBD-intermediate conformation, and the third with two classes in the one-RBD-up conformation. Particles from the second group were combined and subjected to another round of 3D classification using the apex region mask, producing two major classes in the RBD-down conformation, one class in the RBD-intermediate conformation and another one in the one-RBD-up conformation. Particles from the two RBD-down classes were combined with those from the RBD-down class from the first round of signal-subtraction/classification based on the apex region to give a class with total 69,374 particles, which was subjected to 3D autorefinement in C1 symmetry. Additional two rounds of 3D auto-refinement in C3 symmetry using a whole mask of this class, followed by CTF refinement, particle polishing, and 3D auto-refinement, produced a map at 3.1Å resolution. Particles in the two classes of the one-RBD-up conformation from the first round of signal-subtraction/classification based on the apex region were combined with those from the second round of signal-subtraction/classification in the same conformation, and autorefined in C1 symmetry using a whole mask, followed by CTF refinement, particle polishing and a final round of 3D auto-refinement, giving a final reconstruction from 87,330 particles with a map at 3.4Å resolution. The class in the RBD-intermediate conformation from the second round of signal-subtraction/classification based on the apex region was auto-refined in C1 symmetry with a whole mask, producing a map at 4.3Å resolution. Additional rounds of 3D auto-refinement were performed for each class using different sizes of masks at the apex region to improve local resolution near the RBD and NTD. The best density maps were used for model building. At the same time, we have also used cryo-SPARC (*62*) to validate the data processing, which has given similar results.

All resolutions were reported from the gold-standard Fourier shell correlation (FSC) using the 0.143 criterion. Density maps were corrected from the modulation transfer function of the K3 detector and sharpened by applying a temperature factor that was estimated using post-processing in RELION. Local resolution was determined using ResMap (*63*) with half-reconstructions as input maps.

### Model building

The initial templates for model building used our G614 S trimer structures (PDB ID: 7KRQ and PDB ID: 7KRR; ref(*34*)). Several rounds of manual building were performed in Coot (*64*). The model was then refined in Phenix (*65*) against the 3.1Å (RBD-down), 3.4Å (one-RBD-up) cryo-EM maps of the Omicron variant. Iteratively, refinement was performed in both Refmac (real space refinement) and ISOLDE (*66*), and the Phenix refinement strategy included minimization_global, local_grid_search, and adp, with rotamer, Ramachandran, and reference-model restraints, using 7KRQ and 7KRR as the reference model. The refinement statistics are summarized in Table S3. Structural biology applications used in this project were compiled and configured by SBGrid (*67*).

## Acknowledgments

We thank S. Rawson and the SBGrid team for computing support, computing resources from S. Harrison and J. Abraham, K. Arnett for support and advice on the BLI experiments, and S. Harrison and A. Carfi for critical reading of the manuscript. EM data were collected at the Harvard Cryo-EM Center for Structural Biology of Harvard Medical School. We acknowledge support for COVID-19 related structural biology research at Harvard from the Nancy Lurie Marks Family Foundation and the Massachusetts Consortium on Pathogen Readiness (MassCPR). This work was supported by Fast grants by Emergent Ventures (to B.C. and D.R.W.), COVID-19 Award by Massachusetts Consortium on Pathogen Readiness (MassCPR; to B.C. and D.R.W.), and NIH grants AI147884 (to B.C.), AI141002 (to B.C.), AI127193 (to B.C. and James Chou) and AI39538, AI165072, and AI169619 (to D.R.W).

## Author Contribution

B.C., T.X., J.Z., and Y.C. conceived the project. Y.C. expressed and purified the full-length S proteins with help from H.P.. T.X. designed and performed BLI and cell-cell fusion experiments. J.Z. prepared cryo grids and performed EM data collection with contributions from M.L.M., and processed the cryo-EM data, built and refined the atomic models. J.L. and S.W. created the Omicron expression construct and performed the neutralization assays using the MLV-based pseudoviruses. C.L.L. and M.S.S performed the neutralization assays using the HIV-based pseudoviruses. H.Z., K.A. and W.Y. performed the flow cytometry experiments. P.T., A.G. and, D.R.W. produced anti-S monoclonal antibodies. S.R.V. contributed to cell culture and protein production. P.S. provided computational support. All authors analyzed the data. B.C., T.X., J.Z. and Y.C. wrote the manuscript with input from all other authors.

## Competing Interests

W.Y. serves on the scientific advisory boards of Hummingbird Bioscience and GO Therapeutics and is currently an employee of GV20 Therapeutics LLC. All other authors declare no competing interests.

