## Supplemental Figures and Tables for "Structural and functional impact by SARS-CoV-2 Omicron spike mutations"

### Supplementary materials

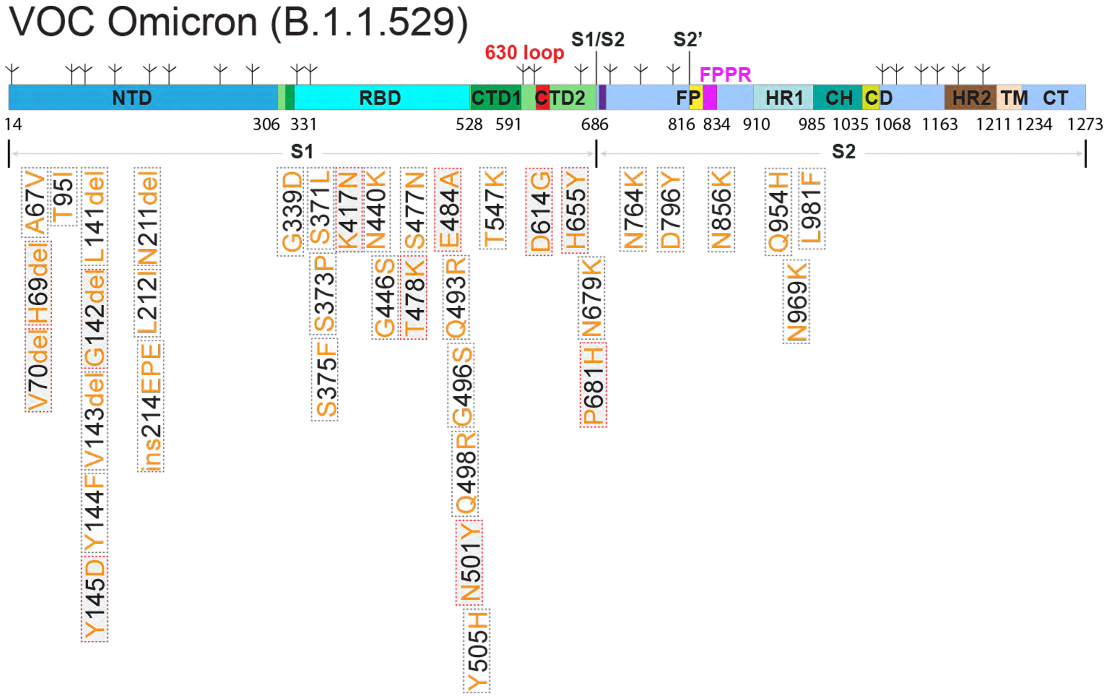

**Figure S1. Schematic representation of full-length Omicron (B.1.1.529) spike (S) protein.** The sequence is derived from an Omicron variant (hCoV-19/South Africa/CERI-KRISP-K032233/2021). Segments of S1 and S2 include: NTD, N-terminal domain; RBD, receptor-binding domain; CTD1, C-terminal domain 1; CTD2, C-terminal domain 2; 630 loop; S1/S2, S1/S2 cleavage site; S2', S2' cleavage site; FP, fusion peptide; FPPR, fusion peptide proximal region; HR1, heptad repeat 1; CH, central helix region; CD, connector domain; HR2, heptad repeat 2; TM, transmembrane anchor; CT, cytoplasmic tail; and tree-like symbols for glycans. Positions of all mutations (from the amino-acid sequence of Wuhan-Hu-1) are indicated and those highlighted in red rectangles are also present in one of the previous VOCs.

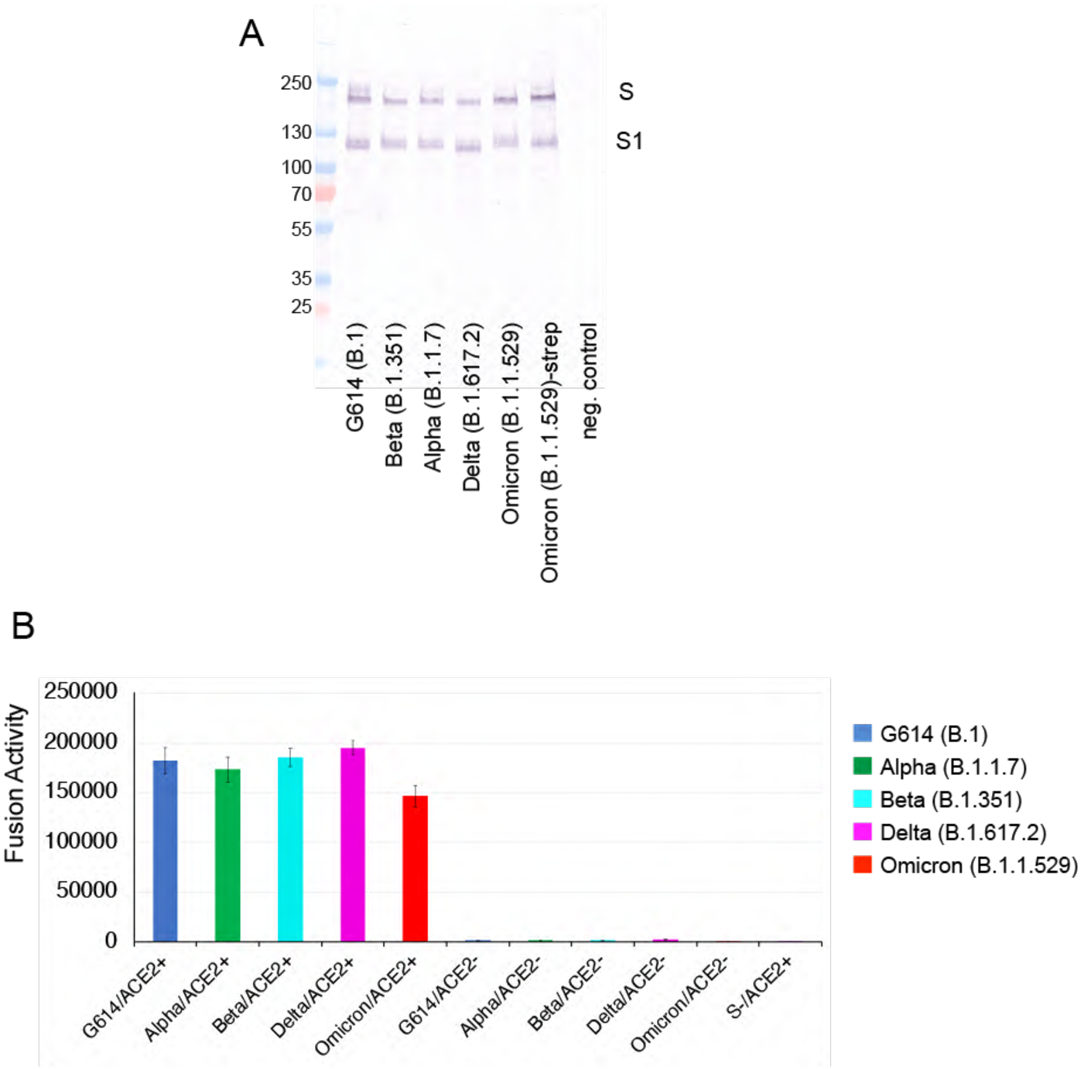

**Figure S2. Expression and cell-cell fusion of the Omicron variant S protein.** (**A**) Expression and processing of the full-length S constructs in HEK293 cells. S protein samples prepared from HEK293 cells transiently transfected with 10 μg of the full-length S expression plasmids were detected by anti-RBD polyclonal antibodies. Bands for the uncleaved S and S1 fragment are indicated. (**B**) HEK293T cells transfected with the untagged, full-length S protein expression plasmids were fused with ACE2-expressing cells. Cell-cell fusion led to reconstitution of α and ω fragments of β-galactosidase yielding an active enzyme, and the fusion activity was then quantified by a chemiluminescent assay. No ACE2 and no S were negative controls.

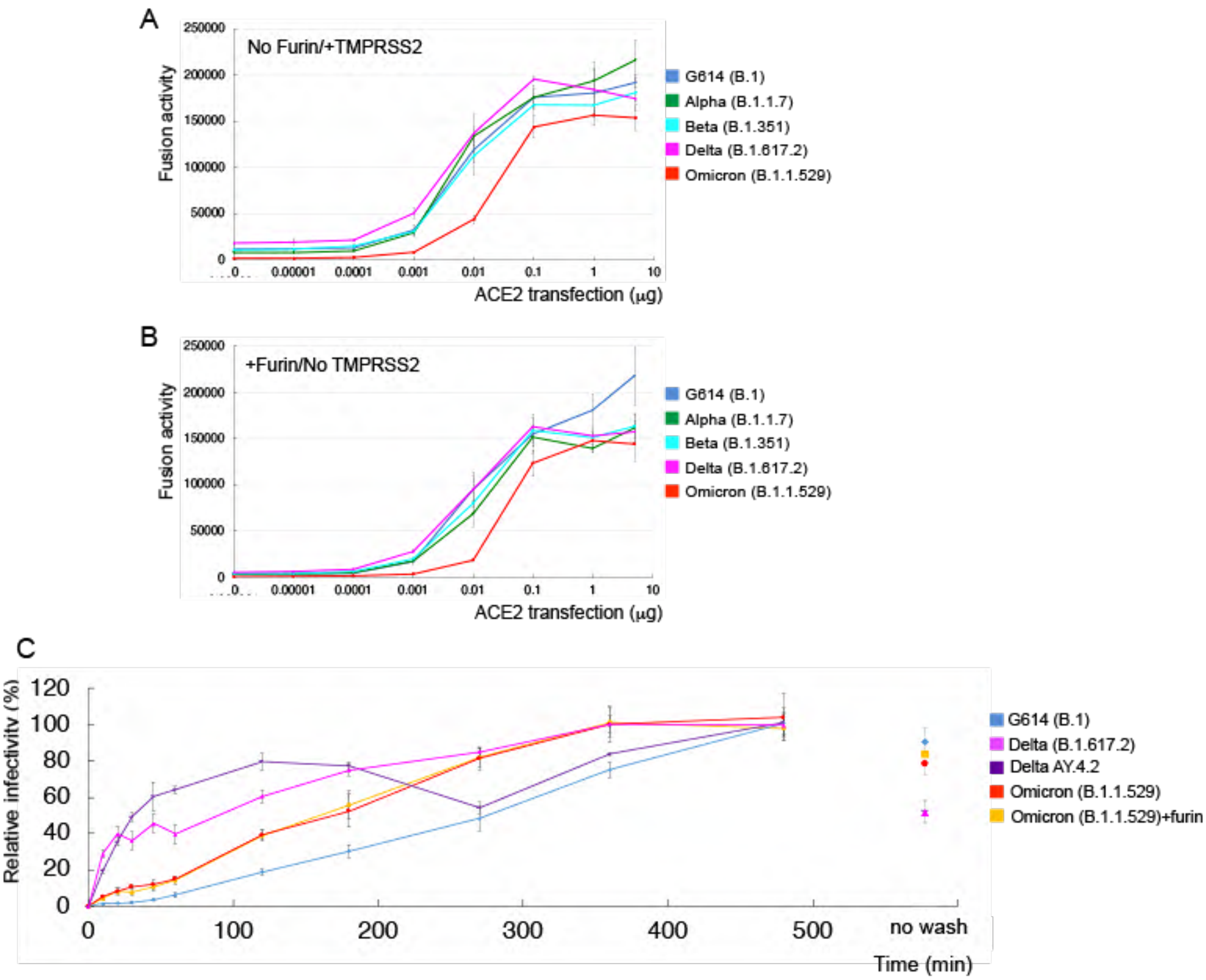

**Figure S3. Comparison of membrane fusion mediated by the Delta variant with that by other variants.** (**A**) Cell-cell fusion mediated by various full-length S proteins with the ACE2-expressing target cells cotransfected with 5 μg TMPRSS2 expression construct. (**B**) Cell-cell fusion mediated by various full-length S proteins expressed in HEK293 cells cotransfected with 5 μg furin expression construct and the ACE2-expressing target cells. (**C**) Time course of infection HEK293-ACE2 cells by MLV-based pseudotyped viruses using various SARS-CoV-2 variant S constructs containing a CT deletion in a single cycle. Infection was initiated by mixing viruses and target cells, and viruses were washed out at each time point as indicated. Delta AY.4.2 is a sublineage of the original Delta (B.1.617.2). Omicron (B.1.1.529)+furin is a virus preparation derived from cotransfection of Omicron S and furin. The experiments were repeated at least two times with independent samples giving similar results.

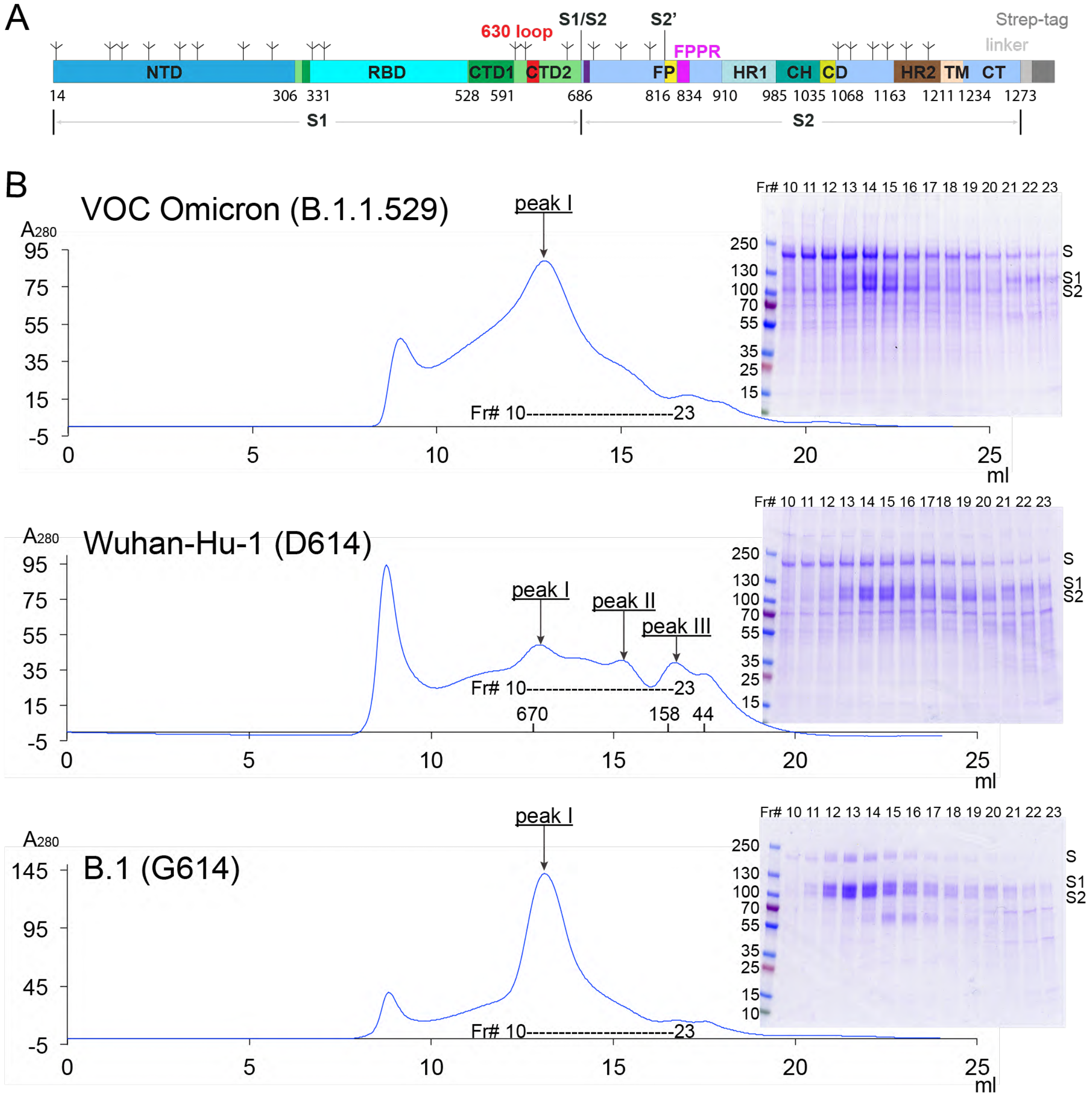

**Figure S4. Production of the full-length S protein from the Omicron variant.** (**A**) A strep-tag for purification was fused to the C-terminus of the full-length S protein by a flexible linker. (**B**) The full-length Omicron S protein was extracted and purified in detergent DDM, and resolved by gel-filtration chromatography on a Superose 6 column. Peak I, the prefusion S trimer; peak II, the postfusion S2 trimer; and peak III, the dissociated monomeric S1. Inset, peak fractions were analyzed by Coomassie stained SDS-PAGE. Labeled bands are S, S1 and S2. Fr#, fraction number. Each experiment was repeated at least three times independently with similar results. The data for the preparations from the Wuhan-Hu-1 (D614) and B.1 (G614), published previously (*34, 36*), are included for convenient comparison.

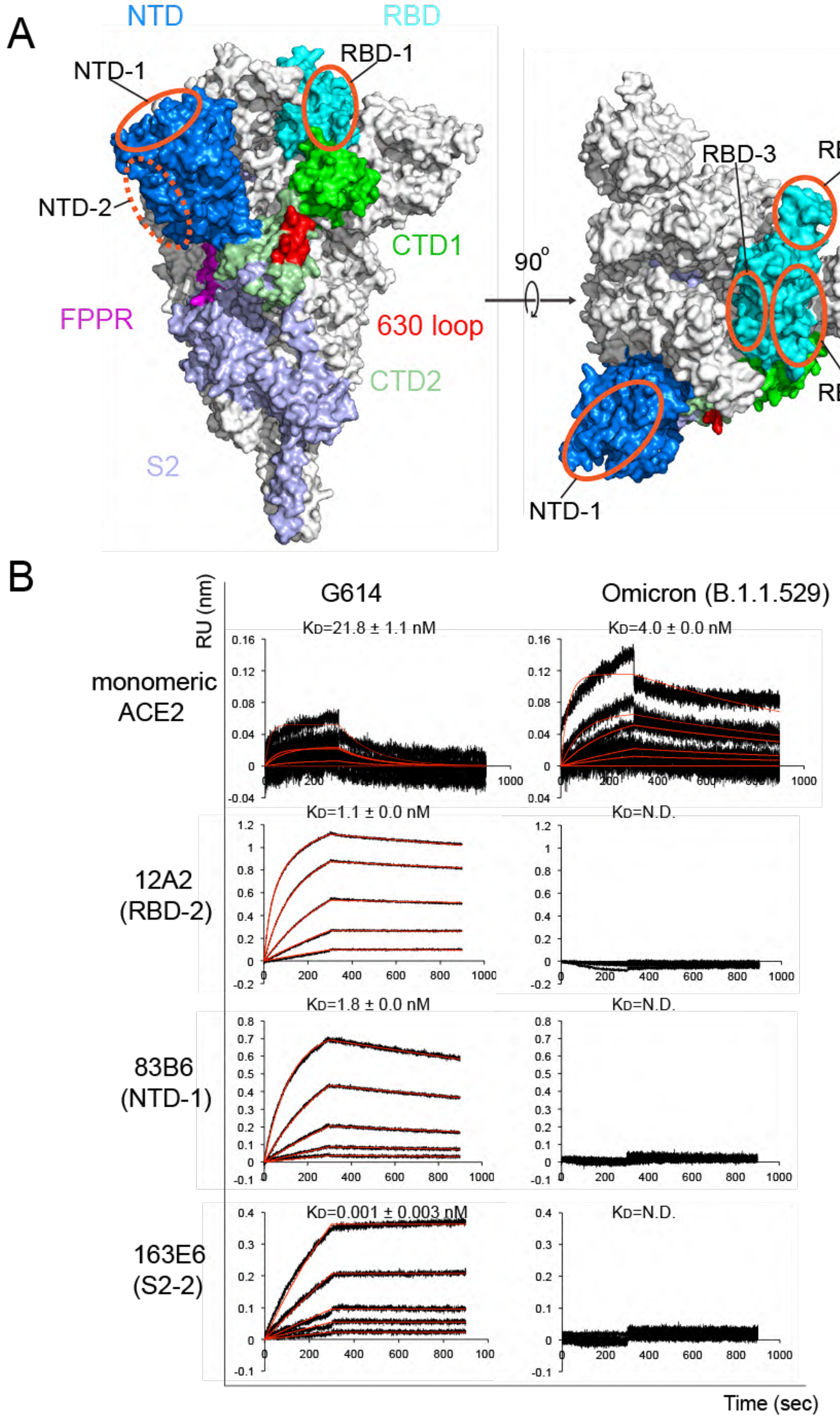
A
NTD
RBD
NTD-1
RBD-1
NTD-2
CTD1
FPPR
630 loop
CTD2
S2
90°
RBD-3
RBD-2
RBD-1
NTD-1
B
RU (nm)
G614
Omicron (B.1.1.529)
monomeric ACE2
KD=21.8 ± 1.1 nM
KD=4.0 ± 0.0 nM
12A2 (RBD-2)
KD=1.1 ± 0.0 nM
KD=N.D.
83B6 (NTD-1)
KD=1.8 ± 0.0 nM
KD=N.D.
163E6 (S2-2)
KD=0.001 ± 0.003 nM
KD=N.D.
Time (sec)

**Figure S5. Additional antigenic properties of the purified full-length Omicron S protein**. (**A**) Antibody competition clusters as described in ref(*39*). Surface regions of the S trimer targeted by antibodies on S1 are highlighted by orange ellipses, including RBD-1, RBD-2, RBD-3, NTD-1 and NTD-2. The exact location of NTD-2 is uncertain and therefore marked with a dashed line. (**B**) Binding analysis of the prefusion S trimers from G614 and Omicron variant with soluble ACE2 constructs was performed by bio-layer interferometry (BLI). For ACE2 binding, purified ACE2 protein was immobilized to AR2G biosensors and dipped into the wells containing each purified S proteins at various concentrations. For antibody binding, various antibodies were immobilized to AHC biosensors and dipped into the wells containing each purified S protein at different concentrations. Binding kinetics were evaluated using a 1:1 Langmuir model except for dimeric ACE2 and antibody G32B6 targeting the RBD-2, which were analyzed by a bivalent binding model. The sensorgrams are in black and the fits in red. Binding constants highlighted by underlines were estimated by steady-state analysis as described in the Methods. RU, response unit. Binding constants are also summarized here and in Table S1. N.D., not determined. All experiments were repeated at least twice with essentially identical results.

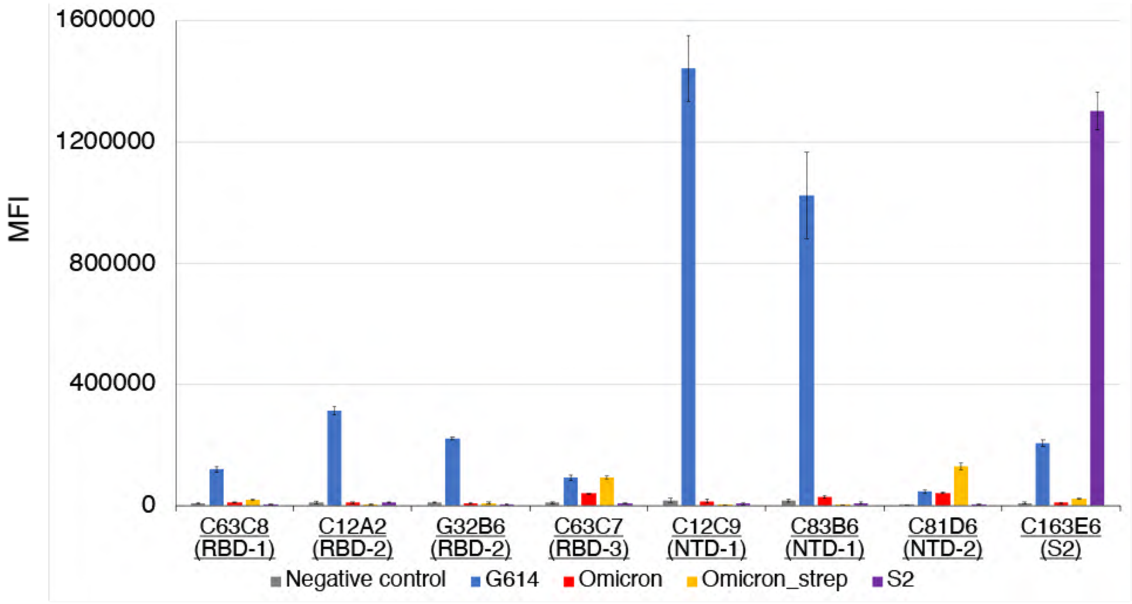

**Figure S6. Antigenic properties of the cell-surface Omicron S protein assessed by flow cytometry.** Antibody binding to the full-length S proteins of the G614 and Omicron variants, as well as an S2 construct expressed on the cell surfaces analyzed by flow cytometry. The antibodies and their targets are indicated. MFI, mean fluorescent intensity. The error bars represent standard errors of mean from measurements using three independently transfected cell samples. The flow cytometry assays were repeated three times with essentially identical results.

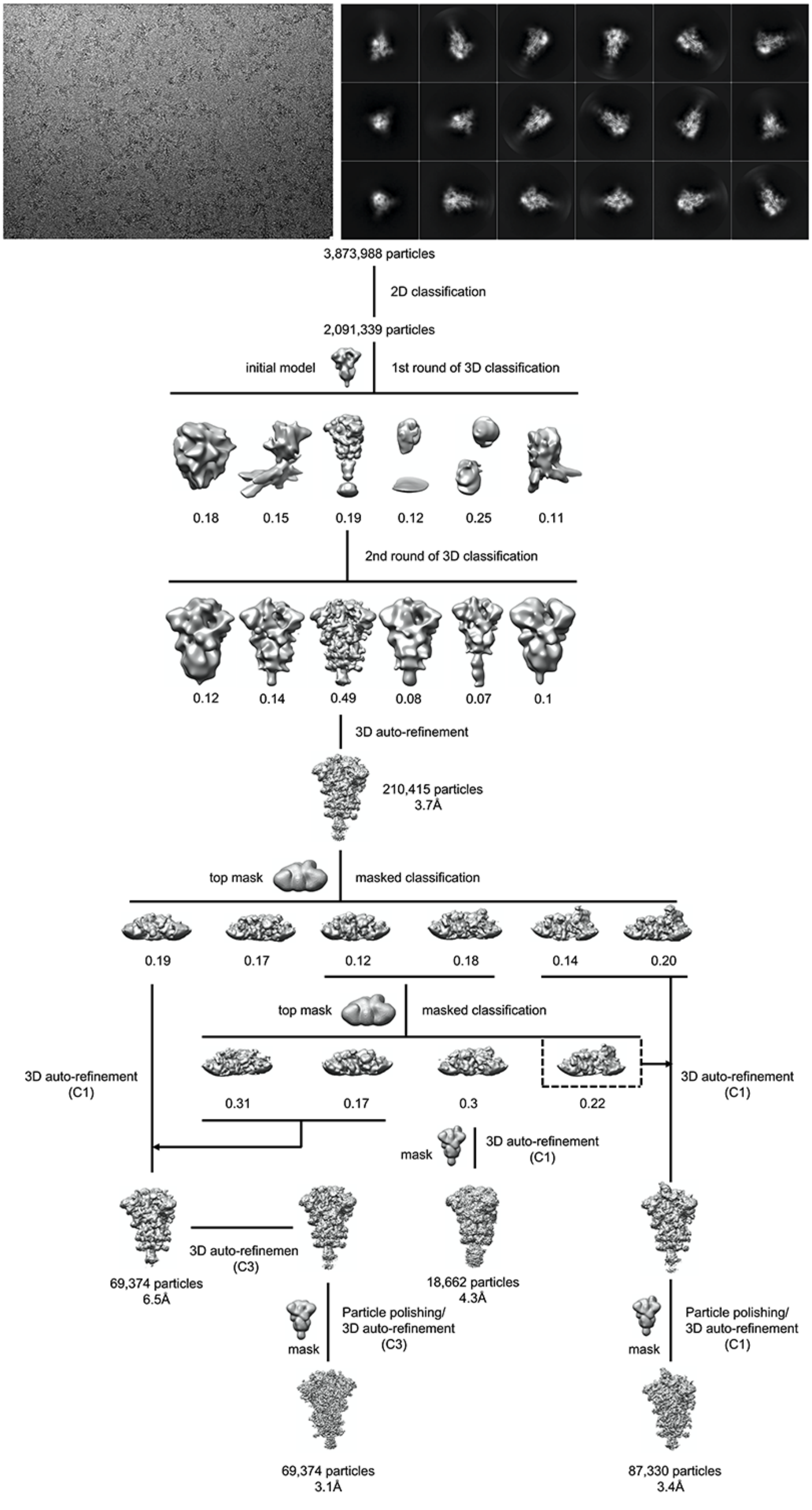
3,873,988 particles
2D classification
2,091,339 particles
initial model
1st round of 3D classification
0.18
0.15
0.19
0.12
0.25
0.11
2nd round of 3D classification
0.12
0.14
0.49
0.08
0.07
0.1
3D auto-refinement
210,415 particles
3.7Å
top mask
masked classification
0.19
0.17
0.12
0.18
0.14
0.20
top mask
masked classification
3D auto-refinement
(C1)
0.31
0.17
0.3
0.22
3D auto-refinement
(C1)
mask
3D auto-refinement
(C1)
3D auto-refinemen
(C3)
69,374 particles
6.5Å
18,662 particles
4.3Å
mask
Particle polishing/
3D auto-refinement
(C3)
mask
Particle polishing/
3D auto-refinement
(C1)
69,374 particles
3.1Å
87,330 particles
3.4Å

**Figure S7. Cryo-EM analysis of the Omicron S trimer.** Top, representative micrograph**,** and 2D averages (box dimension: 396Å) of the cryo-EM particle images of the Omicron S trimer. Bottom, data processing workflow for structure determination.

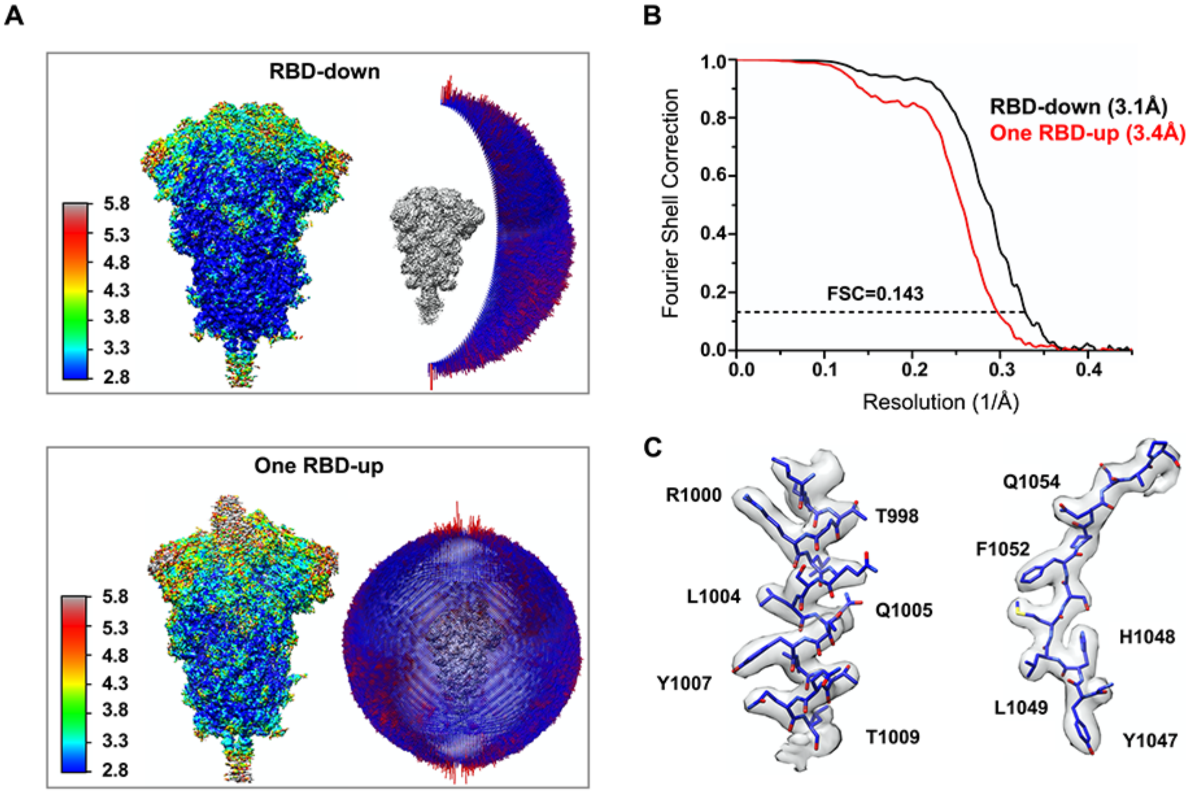

**Figure S8. Analysis of the Omicron S trimer structure.** (**A**) 3D reconstructions of the Omicron trimer preparation in the three-RBD-down and one-RBD-up conformations, respectively, are colored according to local resolution estimated by ResMap. Angular distribution of the cryo-EM particles used in each reconstruction is shown in the side view of the EM map. (**B**) Gold standard FSC curves of the three refined 3D reconstructions of the Omicron S trimer. (**C**) Representative density in gray surface from the EM map of the three-RBD-down conformation.

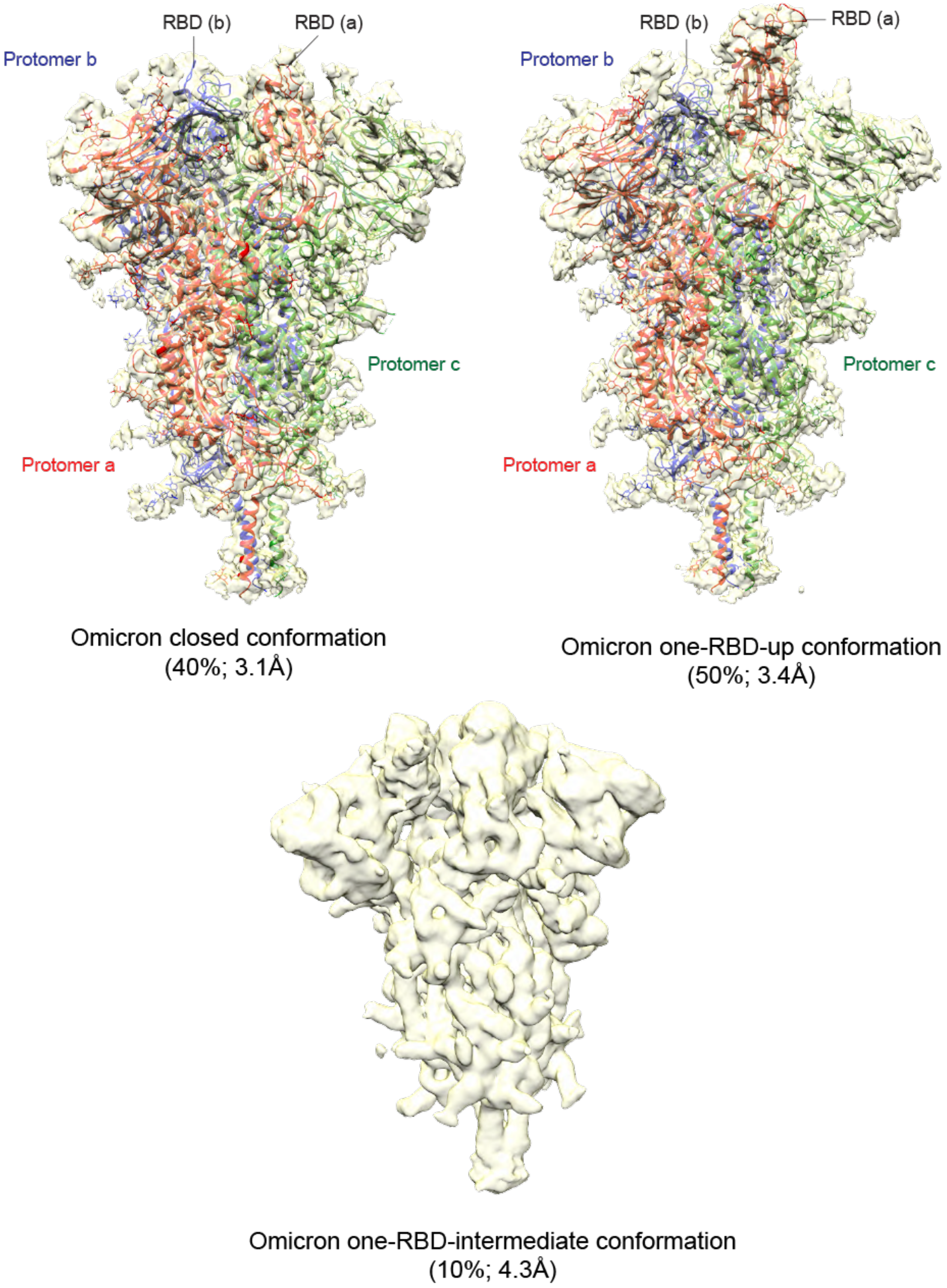

**Figure S9. Cryo-EM structures of the full-length S protein of the Omicron variant**. Three structures of the Omicron S trimer, representing the closed prefusion conformation and one-RBD-up conformations, were modeled based on corresponding cryo-EM density maps at 3.1 and 3.4Å resolution, respectively. Three protomers (a, b, c) are colored in red, blue and green, respectively. RBD locations are indicated. A third class representing a one-RBD-intermediate conformation refined to 4.3Å was not modeled. Particle percentage for each class in the data processing is also indicated, but it may not accurately reflect the conformation distribution of the S trimer in solution.

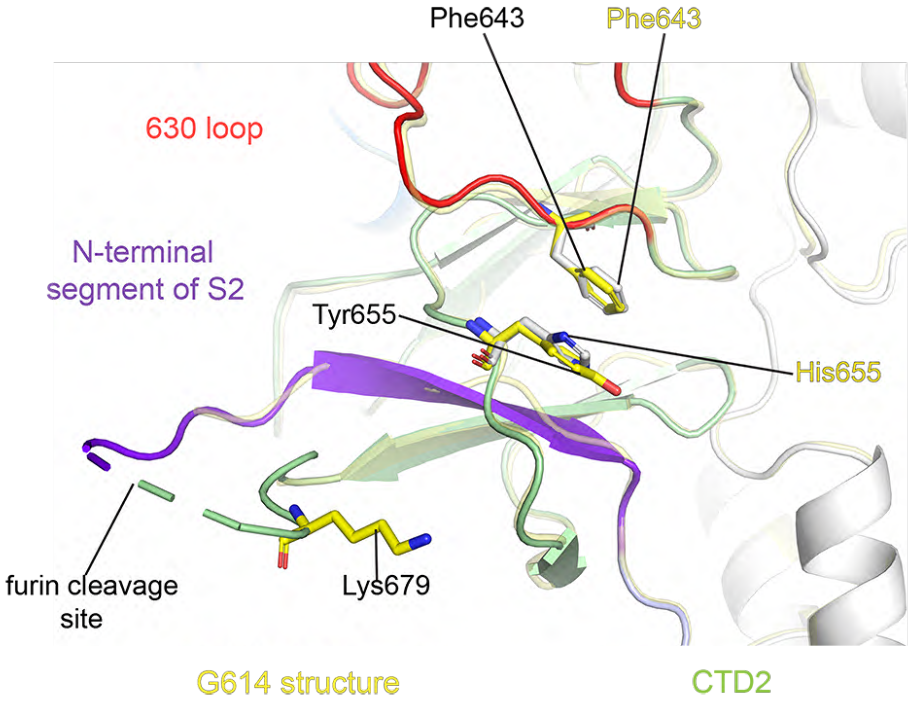

**Figure S10. Additional ordered residues near the furin site in the Omicron structure**. Superposition of the structure of the Omicron S trimer in ribbon representation and various colors with the structure of the G614 S in yellow aligned by S2, showing the region near the furin cleavage site.

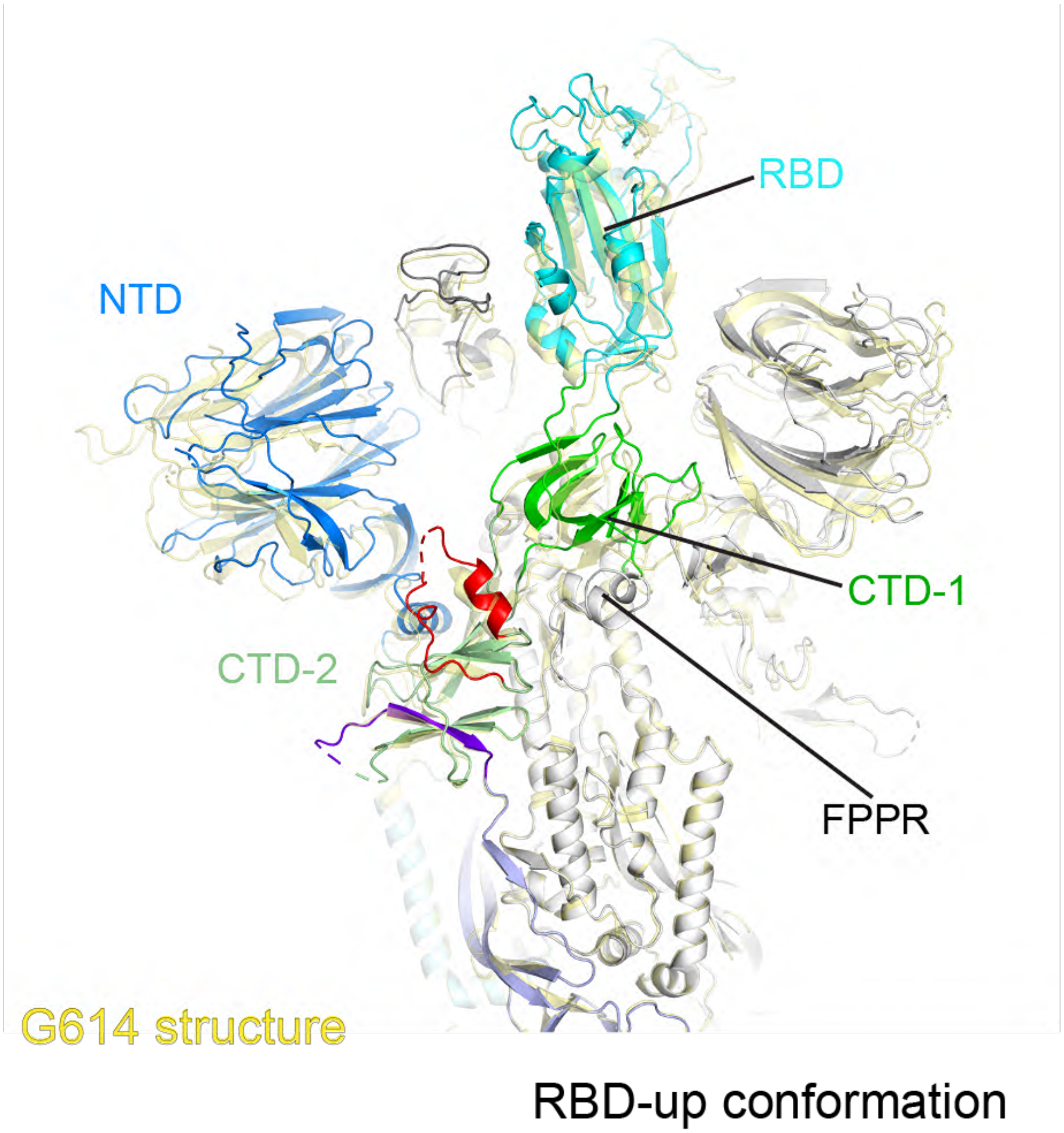

**Figure S11. Superposition of the Omicron and the G614 structures.** Side view of superposition of the one-RBD-up conformation of the Omicron S (various colors) in ribbon diagram, aligned by the S2 portion with the one-RBD-up conformation of the G614 S (yellow). The positions of the RBD in the up conformation and the CTD-1 underneath, as well as the FPPR from the neighboring protomer are indicated.

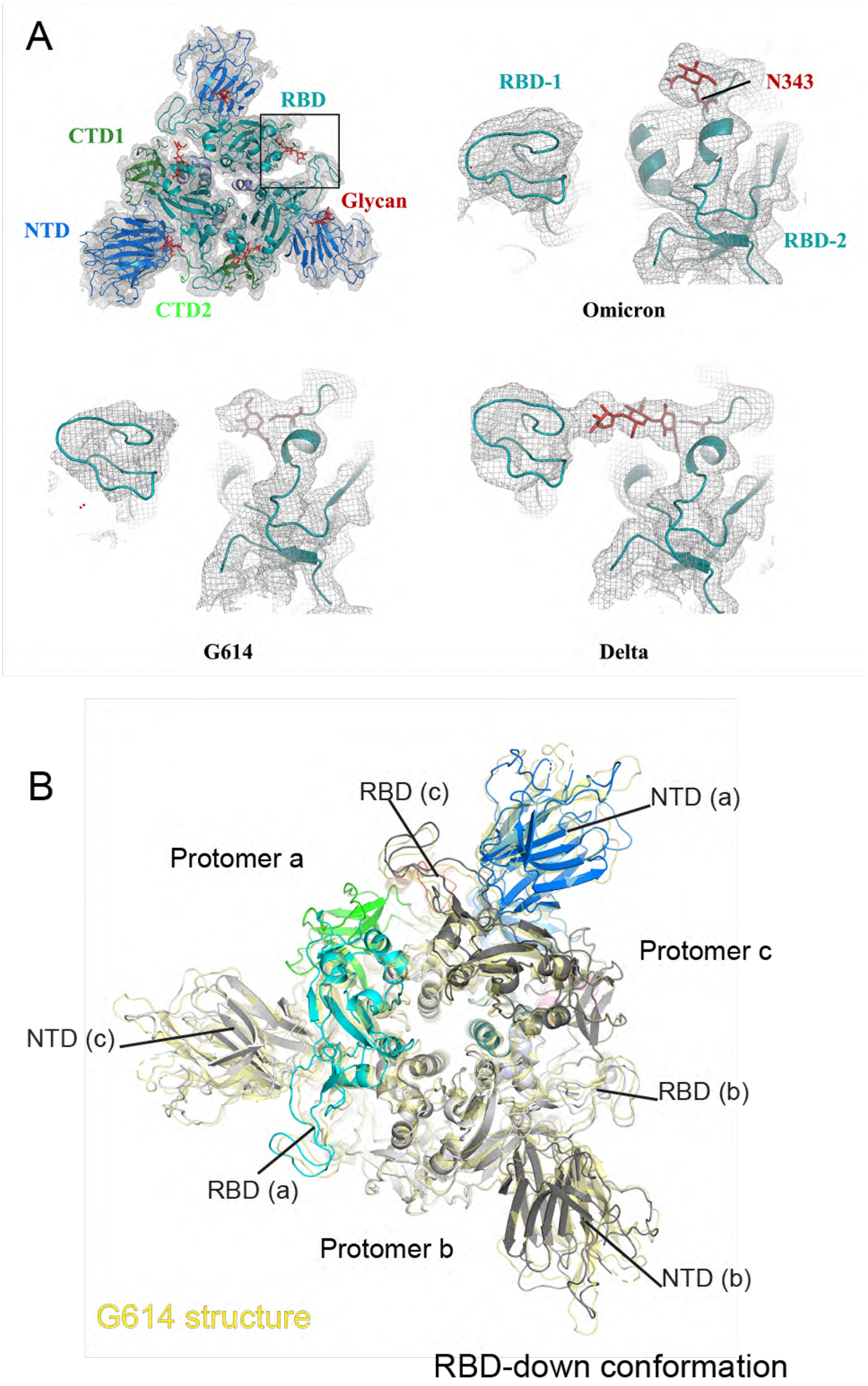

**Figure S12. Comparison of the density of the N-linked glycan at Asn343 and the S1 structure between the Omicron and G614 S trimers.** (**A**) The EM maps of G614, Omicron and Delta trimers in the closed prefusion conformation are compared at the same resolution (3.2Å) and the same contour level. (**B**) Top view of superposition of the closed conformation of the Omicron S (various colors) in ribbon diagram and the G614 S (yellow), aligned by the S2 portion. RBD positions are indicated.

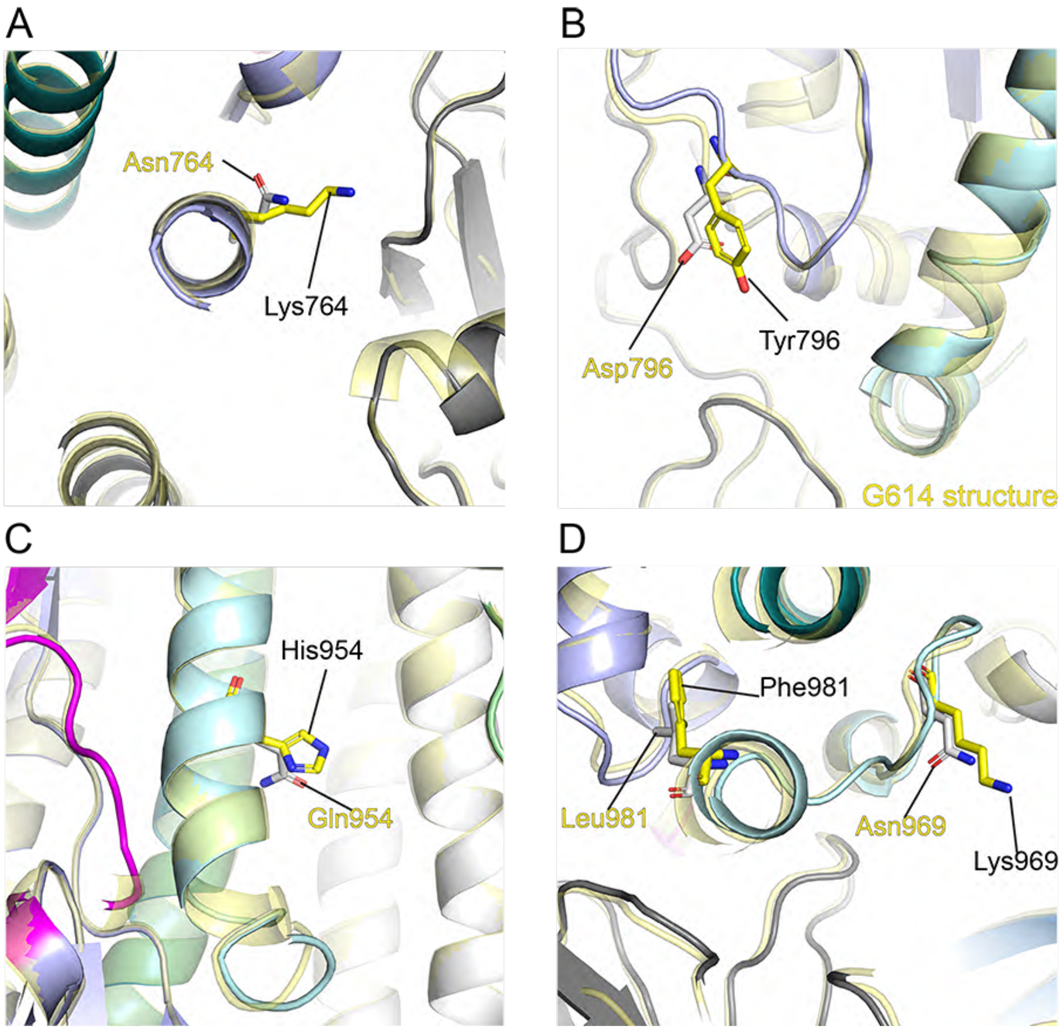

**Figure S13. Additional mutations in the Omicron variants**. (**A**) Superposition of the structure of the Omicron S trimer in ribbon representation and various colors with the structure of the G614 S in yellow aligned by S2, showing the regions near the mutation N764K. (**B**)-(**D**) Similar in (A), the regions near D796Y, Q954H, and N969K/L981F, respectively. All mutations are shown as sticks.

### Table S1. Binding constants of S-ACE2 interaction

| | | $K_D$ (M) | $K_D$ Error | ka (1/Ms) | ka2 | ka Error | ka2 Error | kdis (1/s) | kdis2 | kdis Error | kdis2 Error |
|---|---|---|---|---|---|---|---|---|---|---|---|
| **ACE2-Fc (RBD)** | G614 S | 4.01E-08 | 3.93E-09 | 9.78E+04 | 3.46E+01 | 4.94E+03 | 9.07E-01 | 3.92E-03 | 2.82E-01 | 3.29E-04 | 1.83E-02 |
| | Omicron S | 8.33E-09 | 3.13E-10 | 1.04E+05 | 1.38E+01 | 2.01E+03 | 5.04E+00 | 8.63E-04 | 6.63E-01 | 2.77E-05 | 2.43E-01 |
| **Monomeric ACE2 (RBD)** | G614 S | 2.18E-08 | 1.10E-09 | 3.33E+05 | | 1.61E+04 | | 7.27E-03 | | 1.03E-04 | |
| | Omicron S | 3.97E-09 | 6.00E-11 | 2.19E+05 | | 2.80E+03 | | 8.70E-04 | | 7.05E-06 | |
| **C63C8 (RBD-1)** | G614 S | 3.55E-09 | 2.03E-11 | 1.38E+05 | | 5.91E+02 | | 4.92E-04 | | 1.87E-06 | |
| | Omicron S | N.D. | N.D. | N.D. | | N.D. | | N.D. | | N.D. | |
| **G32B6 (RBD-2)** | G614 S | 5.29E-10 | 8.91E-12 | 2.29E+05 | 3.23E-01 | 1.63E+03 | 3.20E-02 | 1.21E-04 | 3.28E-02 | 1.85E-06 | 3.10E-03 |
| | Omicron S | N.D. | N.D. | N.D. | N.D. | N.D. | N.D. | N.D. | N.D. | N.D. | N.D. |
| **C12A2 (RBD-2)** | G614 S | 1.13E-09 | 2.24E-11 | 1.62E+05 | 1.42E-01 | 1.59E+03 | 1.39E-02 | 1.83E-04 | 2.37E-02 | 3.15E-06 | 2.36E-03 |
| | Omicron S | N.D. | N.D. | N.D. | N.D. | N.D. | N.D. | N.D. | N.D. | N.D. | N.D. |
| **C63C7 (RBD-3)** | G614 S | 4.03E-09 | 1.82E-11 | 1.19E+05 | | 4.13E+02 | | 4.80E-04 | | 1.40E-06 | |
| | Omicron S | 5.63E-09 | 3.05E-11 | 1.06E+05 | | 4.85E+02 | | 5.95E-04 | | 1.72E-06 | |
| **C12C9 (NTD-1)** | G614 S | 5.49E-09 | 4.04E-11 | 6.38E+04 | | 3.88E+02 | | 3.50E-04 | | 1.45E-06 | |
| | Omicron S | N.D. | N.D. | N.D. | | N.D. | | N.D. | | N.D. | |
| **C83B6 (NTD-1)** | G614 S | 1.75E-09 | 5.81E-12 | 1.64E+05 | | 2.91E+02 | | 2.86E-04 | | 8.06E-07 | |
| | Omicron S | N.D. | N.D. | N.D. | | N.D. | | N.D. | | N.D. | |
| **C81D6 (NTD-2)** | G614 S | 7.91E-09 | 4.09E-11 | 1.04E+05 | | 4.81E+02 | | 8.19E-04 | | 1.85E-06 | |
| | Omicron S | 1.62E-09 | 2.00E-11 | 7.23E+04 | | 3.69E+02 | | 1.17E-04 | | 1.32E-06 | |
| **C163E6 (S2-2)** | G614 S | 1.35E-12 | 2.85E-12 | 3.58E+04 | | 3.87E+02 | | <1.0E-07 | | 1.02E-07 | |
| | D Omicron S | N.D. | N.D. | N.D. | | N.D. | | N.D. | | N.D. | |

**Table S2. Neutralization of the SARS-CoV-2 Omicron variant**

| Antibody/ACE2 construct | Neutralization titer (μg/ml) | | | |
|---|---|---|---|---|
| | G614 (B.1) | | Omicron (B.1.1529) | |
| | $IC_{50}$ | $IC_{80}$ | $IC_{50}$ | $IC_{80}$ |
| C63C8 (RBD-1) | 12.578 | 33.721 | >50 | >50 |
| C12A2 (RBD-2) | 0.026 | 0.094 | >50 | >50 |
| G32B6 (RBD-2) | 0.016 | 0.056 | >50 | >50 |
| C63C7 (RBD-3) | 10.494 | >50 | 48.823 | >50 |
| C12C9 (NTD-1) | 0.066 | >50 | >50 | >50 |
| C83B6 (NTD-1) | 0.288 | >50 | >50 | >50 |
| C81D6 (NTD-2) | >50 | >50 | >50 | >50 |
| C163E6 (S2-2) | >50 | >50 | >50 | >50 |
| ACE2-T27W-Fd | 0.098 | 0.468 | 0.036 | 0.126 |
| Positive serum pool_2 (1/x) | 1,172 | 213 | <20 | <20 |
| Normal Human Serum (1/x) | <20 | <20 | <20 | <20 |

### Table S3. Cryo-EM statistics.

| **EM data collection and reconstruction statistics** | | |
|---|---|---|
| Protein | Full-length S of Omicron variant | |
| Microscope | Titan Krios | |
| Voltage(kV) | 300 | |
| Detector | Gatan K3 | |
| Magnification(nominal) | 105,000 | |
| Energy filter slit width (eV) | 20 | |
| Calibrated pixel size (Å/pix) | 0.83 | |
| Exposure rate ($e^-$/pix/sec) | 13.362 | |
| Frames per exposure | 50 | |
| Total electron exposure ($e^-$/Å$^2$) | 53.592 | |
| Exposure per frame ($e^-$/Å$^2$) | 1.072 | |
| Defocus range (μm) | -0.5,-2.2 | |
| Automation software | SerialEM | |
| # of Micrographs used | 34,031 | |
| Particles extracted | 3,873,988 | |
| Particles after 2D classification | 2,091,339 | |
| Class | RBD-down | RBD-up |
| Total # of refined particles | 69,374 | 87,330 |
| Symmetry imposed | C3 | C1 |
| Estimated accuracy of translations/rotations | 0.79/1.78 | 1.35/2.74 |
| Map sharpening B-factor | -94.3 | -105.1 |
| Unmasked Resolution at 0.5/0.143 FSC (Å) | 3.9/3.4 | 4.5/3.8 |
| Masked resolution at 0.5/0.143 FSC (Å) | 3.5/3.1 | 3.9/3.4 |
| Model refinement and validation statistics | | |
| Class | RBD-down | RBD-up |
| PDB | | |
| Composition | | |
| Amino acids | 3357 | 3350 |
| Glycans | 54 | 57 |
| RMSD bonds (Å) | 0.015 | 0.015 |
| RMSD angles (°) | 2.13 | 2.08 |
| Mean B-factors | | |
| Amino acids | 60 | 61 |
| Glycans | 94 | 92 |
| Ramachandran | | |
| Favored (%) | 92.47 | 91.37 |
| Allowed(%) | 6.69 | 7.37 |
| Outliers(%) | 0.84 | 1.26 |
| Rotamer outliers (%) | 4.12 | 3.01 |
| Clash score | 6.47 | 5.80 |
| C-beta outliers (%) | 1.01 | 0.70 |
| CaBLAM outliers (%) | 3.02 | 3.15 |
| CC (mask) | 0.78 | 0.69 |
| CC (volume) | 0.78 | 0.68 |
| MolProbity score | 2.29 | 2.19 |
| EMRinger score | 3.23 | 2.21 |
